# A unifying model for microRNA-guided silencing of messenger RNAs

**DOI:** 10.1101/2025.03.16.643529

**Authors:** Tanmay Chatterjee, Shankar Mandal, Sujay Ray, Alexander Johnson-Buck, Nils G. Walter

## Abstract

Silencing by the miRNA-guided RNA induced silencing complex (miRISC) is dependent on Ago2-chaperoned base pairing between the miRNA 5′ seed (5′S) and a complementary sequence in the 3′ untranslated region of an mRNA. Prevailing mechanistic understanding posits that initial 5′S pairing can further allow functional base pair expansion into the 3′ non-seed (3′NS), while functionally distinct non-canonical pairing was reported between only the 3′NS and the mRNA coding sequence. We developed single-molecule kinetics through equilibrium Poisson sampling (SiMKEPS) to measure highly precise binding and dissociation rate constants of varying-length target sequences to 5′S and 3′NS in a paradigmatic miRISC isolated from human cells, revealing distinct stable states of miRISC with mutually exclusive 5′S and 3′NS pairing. Our data suggest conformational rearrangements of the Ago2-bound miRNA that regulate alternative 5′S-and 3′NS-driven target recognition. The resulting model reconciles previously disparate observations and deepens our acumen for successfully marshaling RNA silencing therapies.

**Graphical abstract:** 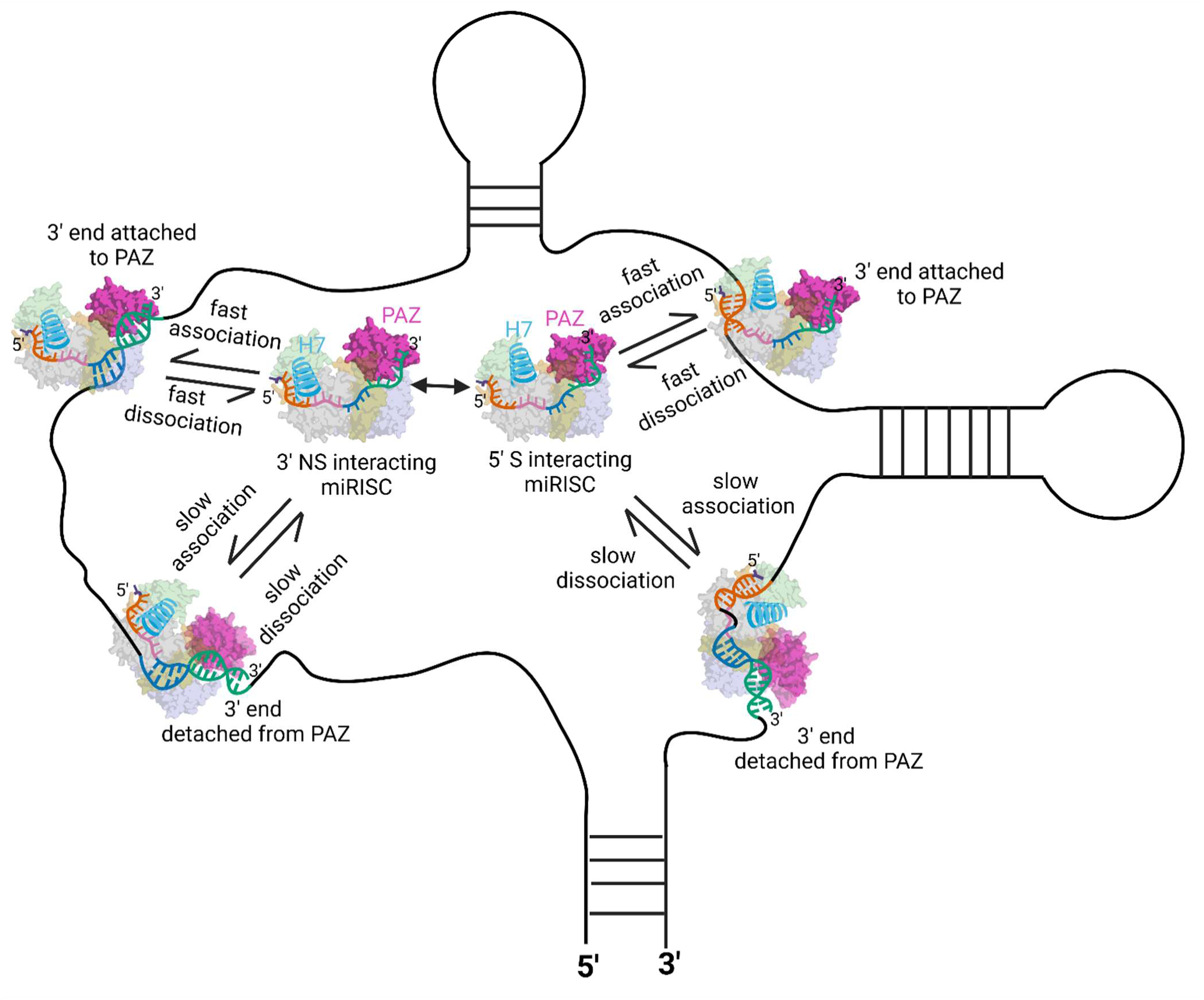

**In brief:** In this study, Chatterjee *et al.* developed a technique to characterize and identify multiple conformations of the RNA-induced silencing complex (RISC) present in human cells. Two distinct pathways of target recognition through different RISC conformations reconcile previously observed divergent findings on the microRNA (miRNA) binding mechanism to messenger RNA (mRNA).

**Highlights:**

- SiMKEPS is developed to precisely measure binding and dissociation of miRISC:mRNA
- Single human miRISC molecules exhibit mutually exclusive seed and non-seed pairing
- SiMKEPS unveils conformational changes of single miRNAs bound to Ago2
- Resulting model reconciles previously divergent findings, supporting RNAi therapies

## INTRODUCTION

MicroRNAs (miRNAs) are a large class of non-protein coding RNAs that play a key role in the regulation of eukaryotic gene expression, under both physiological and pathological conditions, by targeting messenger (m)RNAs for silencing.^1^ Approximately 2,000 distinct miRNAs, some members of sub-families with identical target sequences, regulate an estimated 30% of all human mRNAs.^1–4^ While each miRNA only modestly suppresses a targeted mRNA by first inhibiting translation, then promoting degradation, the promiscuity of each miRNA together with highly combinatorial synergy between miRNAs can lead to rapid changes in the transcriptome and thus gene expression of a human cell, important for proper development and homeostasis.^3–5^ Additionally, many diseases are associated with miRNA dysregulation, from cancer to neurodegenerative and infectious diseases, making an understanding of the detailed targeting mechanism a critical step toward effectively utilizing or sequestering miRNAs for therapeutic applications.^6–9^

Human miRNAs are 21-22 nucleotides in length and associate with Argonaute 2 (Ago2) protein to form miRNA-induced silencing complexes (miRISCs) that engage with the target mRNAs (Figure 1A).^10^

**Figure 1.**
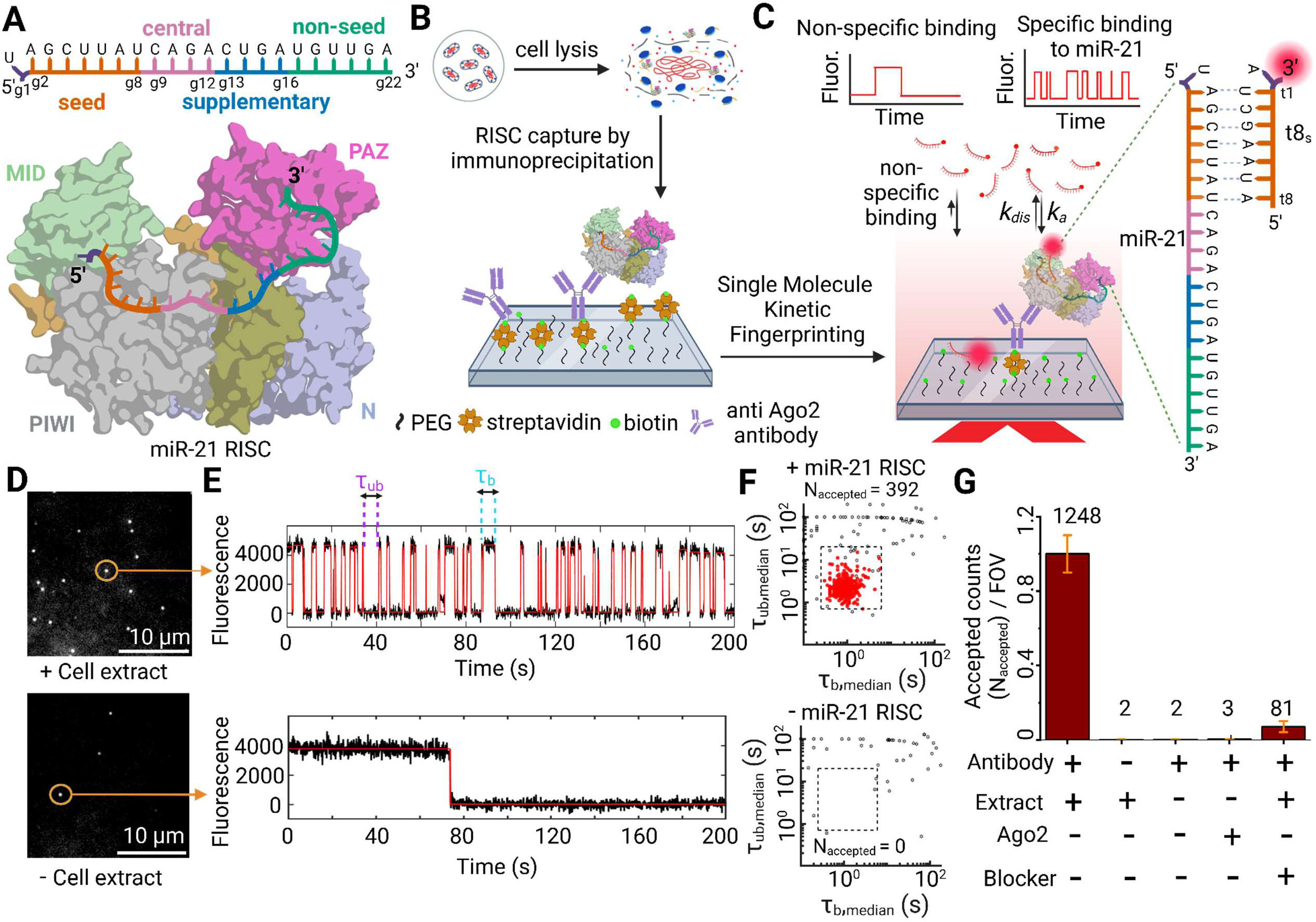
Experimental strategy of single molecule pull-down and analysis of captured RISC. **(A)** Front views of multi-domain (MID, PIWI, N and PAZ domain) human Ago2 protein bound to a guide miR-21. Nucleotides g2 to g8 are called the seed sequence and g13 to g16 are called the supplementary sequence. **(B)** Active RISC was immunoprecipitated from HeLa cell extract and immobilized on a microscope coverslip surface using a biotinylated anti-Ago2 antibody. **(C)** Schematic showing interaction of a kinetic fingerprinting probe with a surface-captured miR-21 RISC yields temporal patterns of repeated binding and dissociation (specific interaction), distinct from nonspecific interaction of the same probes with the surface or other matrix components. **(D)** Single movie frame of a representative portion of a microscope field of view (FOV) showing bright puncta at the locations where single fluorescent probes are bound at or near the imaging surface (upper FOV: with cell extract; Lower FOV: without cell extract) (**E**) The interaction of a Cy5-labeled kinetic fingerprinting probe (t8_7s_) with a surface-captured miR-21 RISC at 21°C yields temporal patterns of repeated binding and dissociation, or kinetic fingerprints (upper trace), distinct from nonspecific interactions of the same probe with the surface or matrix contaminants (lower trace). (**F**) Scatter plots of the median τ_bound_ and τ_unbound_ dwell times for all intensity-versus-time trajectories observed within a single field of view in the presence *(upper)* or absence *(lower)* of miR-21 RISC. Dashed lines indicate thresholds (minimum or maximum) for accepting a trajectory as evidence of a single miR-21 RISC. Black hollow squares represent values from trajectories that do not pass filtering for intensity, signal-to-noise, and/or kinetics, and are not considered to contain sufficient evidence to be classified as binding to miR-21 RISC. Red filled circles represent values of trajectories that pass filtering and are considered positive detection events of single miR-21 RISCs. (**G**) The number of accepted miRISC counts under different conditions. Kinetic filtering keeps the accepted counts nearly zero in absence of either cell extract or capture antibody. Addition of miR-21 inhibitor lowers the accepted counts significantly by blocking access to miR-21 by fluorescent probes. Values shown are the mean of independent duplicates ±SEM.

Nucleotides 2-8 from the 5′-end of the guide miRNA strand (termed g2-g8) are considered the seed sequence that initiates Ago2-chaperoned hybridization with a matching miRNA recognition element (MRE) embedded in the 3′ untranslated region (3′UTR) of the target mRNA.^10–13^ Consistent with this mechanistic model, the Ago2:miRNA complex has been crystallized in a conformation where the 5′ seed (5′S) of the miRNA guide is accessible to bind the target MRE, whereas its 3′ non-seed (3′NS) appears unavailable for target recognition.^10^ Conventional understanding suggests that the 5′S:MRE contact is key for initiating target engagement, which avoids off-target binding by the sequestered 3′NS.^10^ Through a two-step mechanism of target recognition, the 5′S:MRE pairing then induces a conformational change in Ago2—entailing the movement of α-helix-7 and PAZ domain as rigid body—that allows certain miRNAs to expand their base pairing with the target into the 3′NS.^10–13^ In addition to stabilizing miRISC binding to the target, such extended base pairing involving both the 5′S and 3′NS can induce target-directed miRNA degradation (TDMD) for miRNA surveillance.^1,14^

In contrast, genome-wide crosslinking, ligation, and sequencing of hybrids (CLASH) has indicated that 3′NS-only interactions with mRNAs lacking classical 5′S:MRE contacts are also common, with some of these non-canonical interactions functionally validated.^15^ Additionally, recent reports have suggested that miRNAs can target the protein coding sequence (CDS) of an mRNA through such non-canonical interactions associated with a distinct regulatory pathway,^15–18^ challenging conventional models of miRISC action that rely on the 5′S as the sole initiator of any interactions. Heretofore unattainable kinetic profiling of the co-existence, alternation, and crosstalk of miRISC 5′S and 3′NS interactions with a target could elucidate this paradox.

We hypothesized that, intracellularly, miRISC adopts multiple conformations of distinct 5′S and 3′NS accessibility. To test this hypothesis, we combined single molecule pull-down (SiMPull)^19^ of miRISC directly from human cell extract onto a passivated quartz slide with single-molecule kinetic fingerprinting *via* surface-directed total internal reflection fluorescence (TIRF) microscopy^20^ into a technique we term single-molecule kinetics through equilibrium Poisson sampling (SiMKEPS). For SiMKEPS, fluorophore labeled, variable-length RNA probes were chosen as mRNA target mimics, with perfect complementarity to either the 5′S or 3′NS of a miRNA, leading to transient, repeated interactions with the surface-captured, sequence-matched miRISC. Based on hundreds or thousands of individual binding and dissociation events, this approach yielded highly precise binding and dissociation rate constants involving the 5′S and 3′NS of several paradigmatic Ago2-bound miRNAs. After verifying that SiMKEPS distinguishes miRNA sub-family members, even with only subtle distinctions in 5′S sequence, we streamlined our analysis by focusing on hsa-miR-21-5p (miR-21) as one of the most abundant miRNAs in HeLa cells without sub-family members,^4^ but with high relevance for a plethora of human cancers.^21–22^ We discovered two distinct miRISC populations with mutually exclusive 5′S or 3′NS accessibility. In contrast to miR-21 in the absence of Ago2, the interaction kinetics of 5′S and 3′NS of the miRISC-embedded miRNA are indicative of distinct chaperoning by the surrounding protein. In particular, alterations in interaction kinetics indicate that miRISC can undergo a conformational change, likely comprising the movement of the miRNA’s 3′-end relative to the PAZ domain of Ago2, to allow for seed-independent 3′NS pairing with the target. Our findings lead to a model that rationalizes the observed differential, largely mutually exclusive mRNA target engagement by 5′S and 3′NS pairing.

### SiMKEPS detects miRNAs in cellular miRISC with high specificity

To investigate the interaction between the miRISC complex and its target mRNA in the context of translation repression, we designed an *in vitro* assay involving immunoprecipitation of the miRISC directly from HeLa whole cell extract using a validated, highly specific anti-Ago2 antibody,^23^ followed by interrogation of the accessibility of bound miRNAs through iterative hybridization with mRNA mimic probes (see Methods). To validate the pull-down assay, we first immunoprecipitated the miRISC complex (Figure 1A) from the cell extract using the biotinylated anti-Ago2 capture antibody (Figure 1B). A second, orthogonal, fluorophore-labeled anti-Ago2 antibody against a different epitope was used to confirm the specificity of Ago2 pull-down from the cell extract (Figure S1). We found ∼10-15 times more fluorescent spots on the surface only in the presence of both the Ago2 capture antibody and the HeLa cell extract (Figure S1), validating specific capture of endogenous Ago2 and the miRISC complexes it forms.

Next, we investigated the mechanism of interaction between the miRISCs, captured through immunoprecipitated Ago2, and their corresponding mRNA target sequence using SiMKEPS (Figure S2). In SiMKEPS, short fluorescently labeled target mRNA mimics repeatedly interrogate the accessibility of the miRNA in single miRISCs (as shown in Figure 1C), with many complexes subjected to parallel interrogation by widefield TIRF microscopy, to study the mechanisms by which miRISC recognizes and interacts with its target mRNA.

Our study primarily focuses on hsa-miR-21-5p (miR-21), a highly abundant miRNA with ∼10^4^ copies per HeLa cell that lacks a family with significant sequence similarity.^4,24^ This choice simplifies analysis by eliminating the cross-reactive binding we uncovered for miR-16, which shares its seed sequence with family members miR15a and miR195, as well as with members of the miR-103/miR-107 family (Figures S3 and S4). For miR-21, we first employed an 8-nucleotide (nt) fluorophore-labeled MRE-based probe that binds via 7 seed nucleotides, abbreviated as t8_7S_, with perfect complementarity to the seed region g2-g8. This probe, with a predicted melting temperature of 22 °C under our assay conditions, is expected to interact transiently, yet specifically, with surface-immobilized miR-21 RISC. We used TIRF microscopy to detect the fluorescent signal flashes of individual probe binding events and distinguish them from background signal. In the presence of cell extract, the equilibrium binding of the t8_7S_ probe allowed us to detect *N_total_* ∼532 fluorescent spots per movie (2,000 frames, 100 ms exposure, 50 μm x 80 μm field of view=FOV) on the imaging surface (Figure 1D, top panel). Control experiments without cell extract showed considerably fewer fluorescent spots (∼60 fluorescent spots per movie; Figure 1D, bottom panel). In presence of cell extract, the fluorescence intensity versus time trajectories of a majority of individual fluorescent spots show a distinctive kinetic fingerprint that can be used to achieve high discrimination against nonspecific binding to the surface or other matrix components, providing evidence for specific detection of miR-21 RISC (Figure 1E and 1F).

By filtering according to kinetic parameters such as the number of binding and dissociation events (N_b+d_), median dwell time in the bound state (τ_b_, median) and unbound state (τ_ub_, median), we discern true molecular interactions with miR-21 RISC from background signals (Figure 1E-1F, Figure S2; see Methods for details). Specific kinetic parameters were found that yielded an average of *N_accepted_* =1248 ± 150 miR-21 RISC molecules per FOV in experiments with HeLa cell extract (Figure 1G) but only ∼1-2 false positives per FOV in control measurements without cell extract or in the absence of the Ago2-specific capture antibody (Figure 1D, bottom panel). We also verified that the t8_7S_ probe does not interact with purified Ago2 devoid of any loaded miRNAs (*N_accepted_* ∼3 per FOV). Furthermore, blocking the 5′S interaction with a miR-21-specific blocker reduces the number of accepted molecules more than 15-fold, strongly suggesting that the observed kinetic fingerprints following pull-down of Ago2 from cell extract originate from specific interaction of the t8_7S_ probe with miR-21 RISC (Figure 1G).

Although the stochastic behavior of single molecules prevents the precise measurement of rate constants from individual binding events (expected coefficient of variation, CV, of ∼100% for a single-exponential process), the repeated binding of each probe to the same miRISC in SiMKEPS yields many independent measurements (e.g., 30-40 events per molecule over 200 s), greatly enhancing precision. To assess this precision of our single-molecule kinetic analysis, we simulated 400 traces with mean per-molecule rate constants of binding and dissociation equal to those measured for the probe t8_7s_ binding to miR-21 RISC, and fit single-exponential distributions to the bound-and unbound-state lifetimes for each trace separately (see Methods). Measurement error in these simulated traces arises only from: (1) statistical uncertainty due to the finite number of dwell times observed per trace, and (2) fitting error from two-state hidden Markov modeling (HMM) of noisy intensity traces. This allows us to assess precision of the measurement process independent of any molecular heterogeneity that may be present in real samples. The per-molecule estimates of τ_bound_ and τ_unbound_ exhibit well-defined distributions with CVs of 15.8% and 16.8%, respectively, illustrating the precision SiMKEPS achieves through its repeated probing (Figure S5). Although the precision can be increased further through longer observation times (Figure S5), an observation time of 200 s already permits discovery of heterogeneity among single miRISCs. Indeed, when the 392 traces in Figure 1F are subjected to the same single-trace exponential fitting, the estimates of τ_b_ and τ_ub_ show broader distributions with CVs of 58.5% and 67.7%, respectively (Figure S5), indicating substantial trace-to-trace kinetic heterogeneity that we explore further below.

Taken together, these data show that SiMKEPS as a tool provides a robust and precise method for interrogating miRNA within individual miRISC complexes captured directly from cell extracts, with all their inherent heterogeneity, thereby enabling a detailed investigation into miRNA accessibility in an authentic cellular context of target mRNA recognition.

### Mechanistic basis for the conservation of miRISC-mediated target binding

Crystal structure data suggest that mRNA silencing involves a miRISC conformation where the miRNA’s 5′-end is anchored in the Ago2 MID domain and nucleotides g2–g5 are splayed in a helical conformation, rendering it accessible for initial interaction with target mRNA.^10,11^ According to the stepwise mechanism of target recognition, 5’S interaction opens the possibility for supplementary and 3’NS interaction.^10,13,25,26^ However, experimental and computational analyses suggest that there is little advantage of seed plus supplementary interaction over seed interaction for target repression, leading to the conservation of only seed sequences in mammalian target.^27–31^ Though it is known that Argonaute reshapes the binding properties of its nucleic acid guides,^25^ the mechanisms by which Ago2 influences evolutionary conservation of guide nucleotides are not known.

Leveraging our capability of SiMKEPS to precisely measure the dynamic accessibility of miRNA in miRISC *via* many independent binding and dissociation events of single complexes, we investigated the impact of complementarity between the miRNA and the corresponding target mRNA on the identification and recruitment of the miRISC complex to its specific mRNA mimic. To examine how complementarity modulates guide-target interaction, we designed 3′-Cy5-labeled RNA probes of variable length (7 nt to 18 nt, abbreviated as t7_6S_-t18_7S_) as mRNA target mimics to interact with surface-captured miR-21 RISC (Figure 2A). Probes with six or fewer nucleotides complementarity with miR-21 form highly unstable complexes and, as a result, their bound-state lifetimes are too brief to be detected at the highest time resolution of our instrument (20 ms). To obtain as quantitatively precise insights about miRNA-target interactions as possible, we first distinguished the specific interactions of those probes with miR-21 RISC from any non-specific interactions using the SiMKEPS analysis pipeline (Figure S2, S6 and Table S1) and then pursued further analysis of the kinetics. Fitting an exponential cumulative distribution function (CDF) to the cumulative distributions of individual dwell times in the bound state (τ_b_) and unbound state (τ_ub_) obtained from idealized two-state HMM of individual time traces yielded estimates of the association rate constant (*k_a_*), dissociation rate constant (*k_dis_*), and dissociation equilibrium constant (K_D_ = *k_dis_* / *k_a_*) of an interaction (Figures 2B-2D, Figure S7 and Table S2). Fitting a model to distributions of dwell times from ∼5,000–50,000 individual binding and dissociation events per condition provides an accurate estimate of rate constants from SiMKEPS by excluding any influence from nonspecific or background interactions (Figure S7). The 3′-Cy5 label was expected to have minimal effect on the binding kinetics between miRISC and the target since Ago2 does not make any specific contacts with the target downstream of the t1 nucleotide.^10^ Indeed, shifting the position of the Cy5 probe further away from the t1 adenosine does not alter the binding kinetics significantly (Figure S8). Photobleaching has little effect on binding kinetics as the rate constant of dissociation is ∼20 – 2,000 times faster than the rate constant of photobleaching (*k*_photobleach_ ∼ 0.001 s^-1^) under the same measurement conditions (Figure S9).

**Figure 2.**
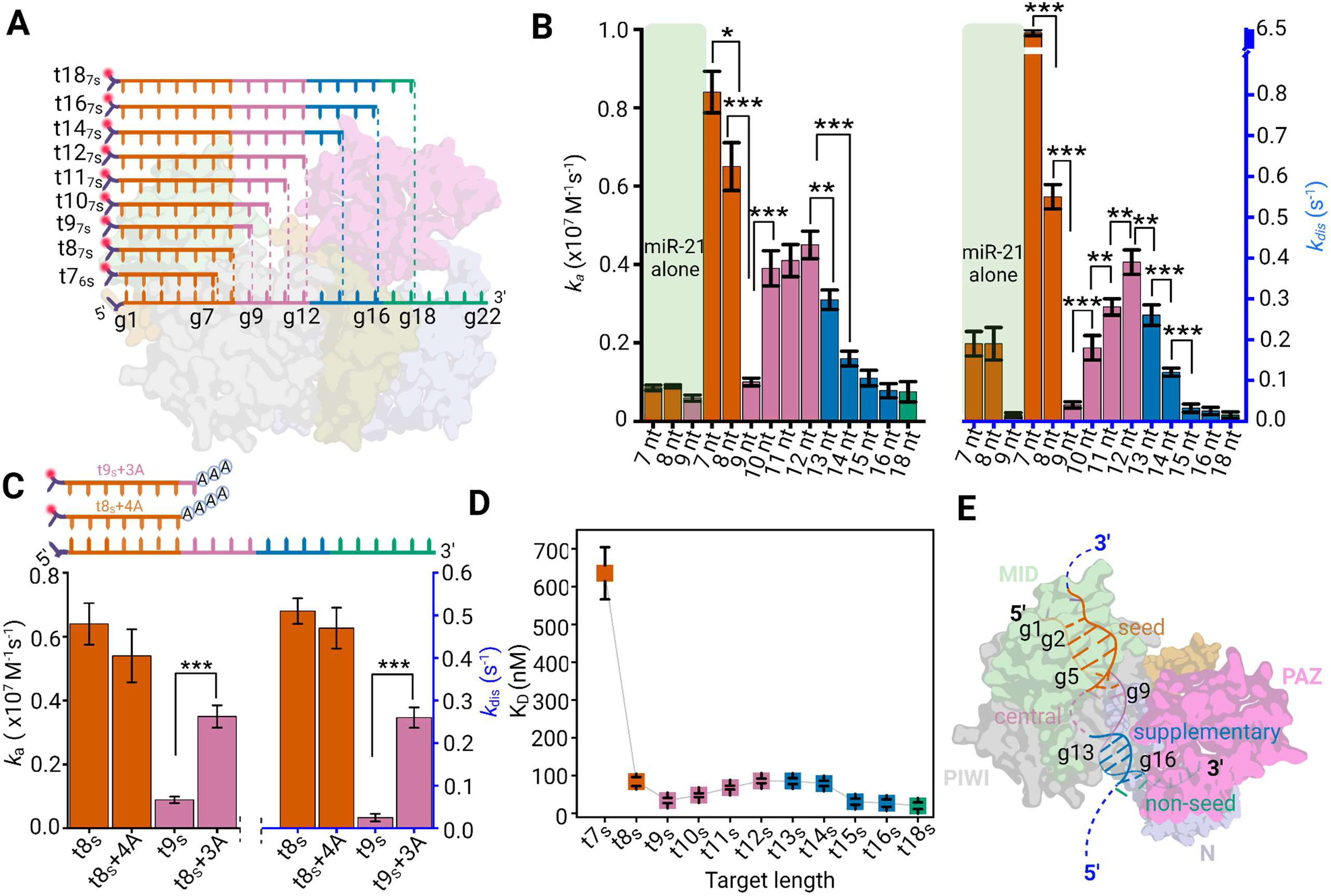
SiMKEPS analysis shows sequence dependence of interactions between miRISC and mRNA target mimics (A) Schematic showing base complementarity within miR-21 RISC ternary complexes with target mimic probes of different lengths. Cy5 probe (shown as red circles) are attached at the 3’-end of each target mimic. **(B)** Comparison of association (*k_a_*) and dissociation (*k_dis_*) rate constants of 7-nt (t7_6S_), 8-nt (t8_7S_) and 9-nt (t9_7s_) probe to miR-21 alone (faint green shaded region). Target sequence-and length-dependent interaction kinetics of the 5′-seed-accessible population of miR-21 RISC at 24° ± 1° C; mean of independent triplicates ± SEM. Statistical significance of differences in the rate constants was determined using two-tailed Student’s test (t test). P values < 0.1 were considered significant (***p < 0.01, **p < 0.05, *p < 0.1). (**C**) Dependence of miRNA-target mimic interaction kinetics on additional unpaired nucleotides (multiple adenines) at the 5′-end of t8_7s_ and t9_s_, showing that the impact of additional nucleotides is dependent on the presence or absence of g9:t9 base pairing in the ternary complex. (**D**) Binding affinity (dissociation equilibrium constant, K_D_) of target mimics having varying degrees of guide complementarity to miR-21 RISC. Mean of independent triplicates ± SEM. (**F**) Model of the participation of nucleotides from the guide strand’s seed (g2–g8) and supplementary (g13–g16) regions in increasing the binding affinity of the miR-21 RISC for the target RNA. Guide-target complementarity in the central (g9–g13) and 3′-end non-seed region (g17– g18) does not correlate with the overall stability of the ternary complex, consistent with the typical lack of base pairing at the central and 3′ non-seed region in this miRISC conformation.

miR-21 RISCs show an ∼11-fold faster association rate constant with t7_6s_ (0.89 x10^7^ M^-1^s^-1^ at 24 °C) than to protein-free miR-21 alone (0.75 x10^6^ M^-1^s^-1^ at 24 °C), consistent with the expectation that Ago2 pre-organizes the 5′S region of the miRNA for faster association with the target (Figure 2B and Figure S10).^10^ Furthermore, there is a considerable increase in target affinity (∼8-fold reduction in K_D_) for miR-21 RISC when extending the mRNA mimic by just one nucleotide, from t7_6s_ to t8_7s_ (Figure 2B and 2D). In contrast, the association rate constant, dissociation rate constant and target affinity of miR-21 alone do not change significantly between the same pair of probes (∼1.2-fold reduction in K_D_; Figure 2B), perhaps because the additional base pair in t8_7s_ is a terminal A-U base pair that may not stably form at room temperature in the case of a protein-free miR-21 that lacks the pre-organization by Ago2. Our data thus suggest that miRISC organizes the nucleotides in the 5’S region such that target binding is stabilized only upon formation of at least 7 base pairs between the target and 5′S (g2–g8), consistent with previous observations.^11^ Indeed, one study has shown that a 6-mer seed match was insufficient for miRNA repression even when multiple copies were inserted in the 3′ UTR of mRNAs, whereas a single insertion rendered the resulting 7-mer seed sequence sufficient for gene silencing.^27^

Interestingly, the presence of adenosine at the t1 position (which is complementary to the g1 nucleotide of the guide miRNA) does not alter the association rate constant significantly, implying that g1 is not involved in the initial target recognition. However, the presence of adenosine at position t1 does increase affinity by approximately 2-fold and 4-fold compared to targets with cytosine or thymine at t1, respectively, primarily by lowering the dissociation rate constant (Figure S11). We can rationalize this relative strength of the interactions since it is known that an adenosine at t1 enhances, once the ternary complex is established, the stability of the interaction with the guide miRNA by forming hydrogen bonds between Ser^561^ and the N6 amine of adenine (A).^10^ In contrast to thymine (T) or uracil (U), cytosine (C) at t1 can conceivably stabilize the interaction because it can form a single hydrogen bond between its N4 amine and Ser^561^ (Figure S11). Still, since the structure of C as a pyrimidine differs from that of A as a purine, it is expected to result in a weaker hydrogen bond and a correspondingly lower stabilization of the complex. Instead, the carbonyl group of guanine (G) can accept a hydrogen bond from the hydroxyl group of Ser^561^, offering stabilization that becomes comparable to that provided by C. Neither U nor T possess the N6 amine of A or N4 amine of C, and their structural mismatch with the purines leads to further destabilization when either is present at the t1 position. Finally, the presence of a methyl group in thymine is expected to further destabilize the resulting ternary complex compared to uracil (Figure S11).

Long-lived binding of miRISC to target mRNAs is crucial for effective miRNA-mediated translational repression.^11,25^ However, in vertebrates target complementarity is generally only conserved up to the eighth nucleotide of the guide miRNA (position: g1–g8). We thus hypothesized that contiguous target complementarity extending beyond the g8 position threshold does not confer any additional affinity to the ternary complex. To test this hypothesis, we studied binding of the seed probes t7_6S_–t18_7S_ to miR-21 RISC. We found that, while the dissociation rate constant of t9_7S_ is ∼12 times slower (*k*_dis_ ∼0.04 s^-1^) than t8_7S_ (*k*_dis_ ∼0.51 s^-1^) its association rate constant (*k*_a_ ∼0.11 x 10^7^ M^-1^s^-1^) is ∼6 times slower than that of t8_7S_ (*k*_a_ ∼0.65 x 10^7^ M^-1^s^-1^). Thus, for a short target, contiguous base pairing up to 9 nt (position: g1–g9) only modestly increases the affinity (∼2-fold) compared to the contiguous 8 nt base pairing (position: g1– g8). Contrary to expectations from hybridization thermodynamics, extending target complementarity into the central region (t10_10S_ to t12_7S_) gradually *destabilizes* the ternary complex, increasing *k_a_* but also raising *k_dis_* by a slightly larger factor (Figure 2B), resulting in a net increase in *K_D_* (Figure 2D).

We hypothesized that the much longer target mRNA in physiological contexts may result in unfavorable interactions with the Ago2 central cleft when the g9:t9 base pair is established, negating any stabilizing effects of complementarity beyond the eighth nucleotide (g8). To test this hypothesis, we designed target mimics extended at the 5′-end with 4 or 3 adenosine nucleotides (t8_7S_+4A and t9_7S_+3A, respectively), chosen to not be complementary to the guide miR-21 sequence (Figure 2C). As expected, we observed a significant decrease in target affinity for t9_7S_+3A, but not t8_7S_+4A, which raises the dissociation rate constant back to similar levels as observed for t8_7S_ (Figure 2D and Figure S12). We conclude that base pairing at the g9:t9 position of an mRNA-miRISC ternary complex is hindered for a longer target like mRNA, likely due to steric clashes of nucleobases t10–t12 with the central region of Ago2. The result is consistent with the observation that the t9 nucleotide was disordered in crystals containing a longer target like mRNA.^10^

After the central region, the gradual addition of nucleotides into the supplementary region (corresponding to guide nucleotides g13–g16) stabilizes the ternary complex, albeit incrementally (Figure 2B). In particular, extending target length to sixteen nucleotides (t16_7S_), which ensures complete complementarity across the seed (g2–g8), central (g19–g12), and supplementary regions (g13–g16), leads to a ∼30-fold reduction in dissociation rate constant when compared to a target that only pairs with the seed region (t8_7S_, Figure 2B). Intriguingly, the association rate constant of t16_7S_ decreases ∼10-fold compared to that of t8_7S_ (Figure 2B and Table S2), thus modestly increasing the affinity (K_D_ (t16_7S_)/ K_D_ (t8_7S_) ∼ 3). This observation is consistent with previous report of an ∼10 times slower association rate constant for a fully complementary target than that for a seed matching target.^28^ Notably, additional complementary bases beyond g16 of the guide miRNA do not provide any further increase in affinity, possibly due to the passing of the 3’NS tail (g17 – g21) through a narrow gap between the N and PAZ domains that hinders effective base pairing.

Based on these SiMKEPS findings, we propose a conceptual framework of how miRISC identifies its target in a 5’S-binding conformation. We posit that target complementarity to g2–g8 of the guide miRNA is crucial and sufficient for the initial fast and stable target interaction, whereas contacts with nucleotides outside of the seed region (g2–g8) considerably slow the association rate constant, overall only modestly increasing the affinity (∼3-fold) (Figure 2D). The slow association rate constant for a target interacting with guide nucleotides outside the seed region (g2–g8) are consistent with previous reports suggesting that these interactions require an altered conformation of miRISC^10,26^, involving the movement of the 3’-end in the PAZ domain in a way that slows association with the seed region. Consequently, guide-target complementarity in the central region (g9–g12) and the 3′-end non-seed region (g17–g18) does not contribute to the stability of the ternary complex (Figure 2D and 2E).

Comparative analysis in flies reveals that most sites under selection do not have more 3′-supplementary pairing than what would occur by chance.^29^ Similarly, parallel analyses in mammals have led to the dismissal of 3′-supplementary pairing altogether.^30^ In fact, pairing and energy-based criteria originally intended to identify and rank 3′-supplementary pairing have proven to lack predictive value.^31^ Our experimental results demonstrate why this is the case with the mammalian miRISC. For 5′S-accessible miRISC conformations, 8 nt (g1–g8) seed complementarity is both necessary and sufficient for miRISC function, while target complementarity beyond the canonical seed site (g2–g8) does not offer enough of an increase in binding affinity to be conserved in target mRNA sequences. The modest contribution of the supplementary base pairing to target affinity can explain why in mammals less than 5% of evolutionarily conserved, predicted miRNA-binding sites include conserved 3′ pairing.^1^ However, targets with very weak or partial seed pairing can still gain affinity from supplementary interactions, or supplementary interaction can fine-tune the guide-target interaction, especially when the supplementary bases have high GC content.

### Stable 5’-seeded target interactions are intermittently inhibited in miRISC

Our single trace fitting and simulations suggest heterogeneity in the interaction kinetics between miRISC and the target mimic t8_7s_. To further investigate the nature of this heterogeneity, we performed conventional fitting of cumulative dwell time distributions; these distributions showed bi-exponential behavior, further indicating heterogeneity in the guide-target interaction (Figure S13). Further, scatter plots of the median per-molecule τ*_bound_* and τ*_unbound_* dwell times exhibit broad distributions compared to the simulated scatter plots, suggestive of heterogeneous kinetic behavior (Figure 3A), which was corroborated by per-molecule fitting of dwell time distributions (Figure S5). Inspection of individual target mimic binding trajectories revealed that frequent seed binding events are separated by periods of more sporadic or almost no seed binding (Figure 3B). A fraction of miRISC displays at least one interval when no target binding is observed, with varying durations (Figure 3C). A subset of miR-21 RISC complexes (∼30%, blue bars in the histogram of Figure 3C) exhibit at least one period of 5′S inaccessibility to the target mimic lasting ≥30 s, ∼80-fold longer than the τ_unbound_ in 50 nM target mimic and thus expected to occur with negligible probability in a homogeneous population (Figures 3C). Furthermore, a significant fraction of traces exhibits even longer individual unbound dwell times.

**Figure 3.**
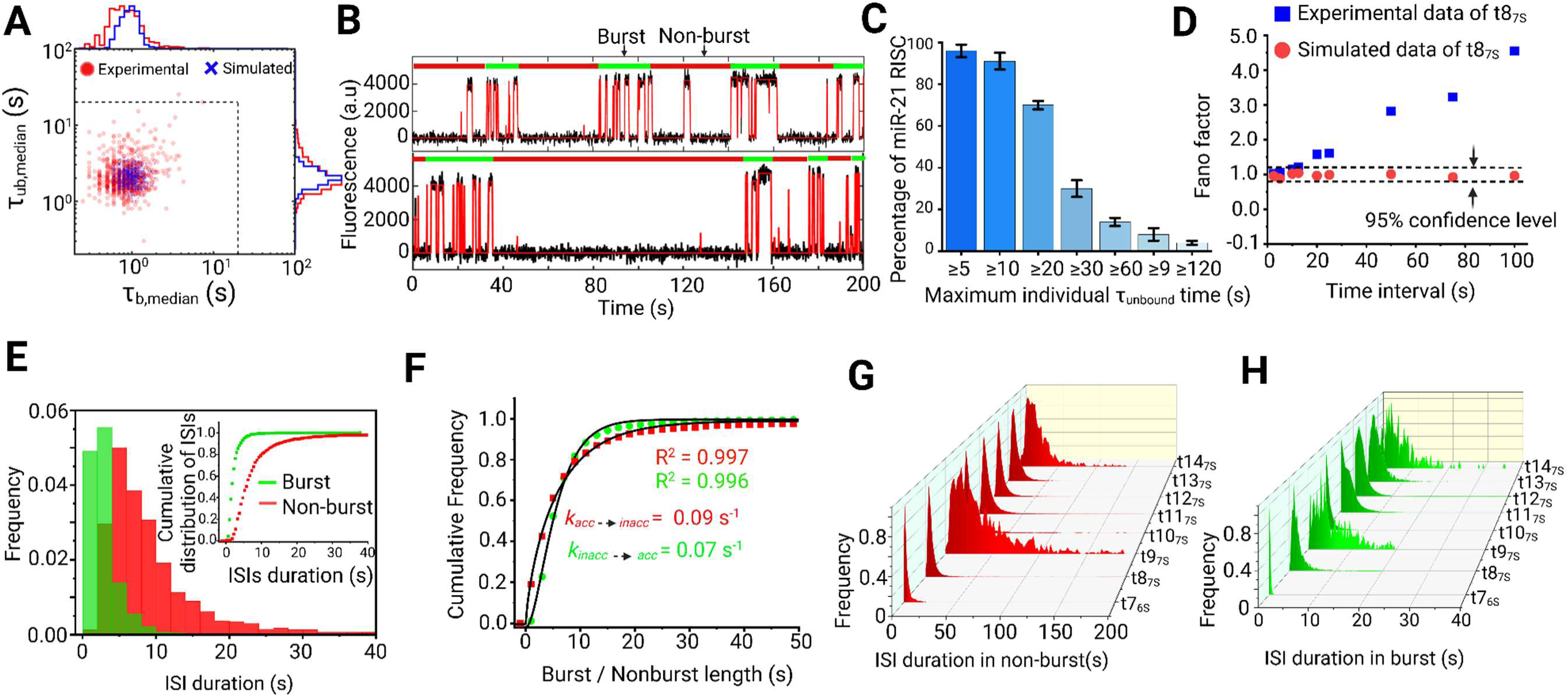
Single miRISC complexes are characterized by burst behavior, with periods of higher and lower seed region accessibility. **(A)** Median dwell time correlation plot for experimental (red circle) and simulated (blue cross) data set for the interaction of miR-21 RISC and t8_7S_ target mimic (number of molecules considered *N* = 321). **(B)** Two representative intensity-versus-time trajectories for the interaction of miR-21 RISC with the t8_7S_ probe with burst (green bars) and non-burst (red bars) periods of target binding detected through spike train analysis. **(C)** Bar plot showing the percentage of miR-21 RISC complexes with at least one variable maximum individual τ*_unbound_* dwell time. Mean of independent triplicates ± SEM. **(D)** The Fano factor calculated across various time intervals for the interaction of miR-21 RISC and t8_7S_ target mimic indicates non-Poissonian statistics (blue squares). The Fano factor confidence interval for Poisson behavior. **(E)** Cumulative histogram displaying the distribution of interspike intervals (ISIs) during burst (green) and non-burst (red) periods. **(F)** Fitting of the burst and non-burst time cumulative distribution function with a single-exponential equation gives the rate constant of conformational transition of miRISC having differential seed accessibility. (**G**) Guide-target complementarity dependent variation of inter spike intervals in non-burst period. (**H**) Guide-target complementarity dependent variation of inter spike intervals in burst period.

The target-mimic-binding events observed here for miRISC strongly resemble neuronal spike trains, where neuronal firing is detected as a sudden, transient increase (spikes) in electrical activity in response to external stimuli.^32–34^ Such spike trains comprise short intervals of high firing activity, called ‘bursts,’ and periods of low or no firing activity, called ‘non-bursts.’ We tested this resemblance by calculating the Fano factor, defined as the variance of the number of spikes within a certain time frame divided by the mean number.^32^ For a random Poisson process, the Fano factor is equal to one regardless of the choice of time interval.^34^ Simulated traces characterized by a single set of rate constants do not deviate from ideal Poisson behavior and the calculated Fano factor remains within the bounds of the 95% confidence interval of a Poissonian process, indicating random Poisson statistics (Figure 3D). In contrast, the binding kinetics of the anti-seed target mimic t8_7S_ to miR-21 RISC clearly deviate from ideal Poisson behavior with a calculated Fano factor much greater than one and increasing at longer time intervals (Figure 3D). The most likely explanation for this heterogeneity is that periods of high seed accessibility are followed by periods of low or no accessibility, and vice versa. Therefore, we applied the Rank Surprise (RS) method of burst detection to detect regions of high spike activity. When the duration of inter-spike intervals (ISIs), equivalent to τ*_unbound_* dwell times, were plotted from the bursting and non-bursting periods, two distinct intramolecular behaviors were observed (Figure 3E). A single miRISC typically interconverts from periods of high seed accessibility to those of low or no seed accessibility. The ISIs in both burst and non-burst periods show the narrowest distribution for t7_6S_ but gradually broaden, reaching their maxima for t9_7S_. After t9_7S_, ISIs in both burst and non-burst periods again become narrower for target lengths up to t12_7S_, then broaden once more for t13_7S_ and t14_7S_, where contacts with the supplementary bases are established and begin stabilizing the guide-target interaction (Figure 2B, 3G and 3H).

To rule out the possibility that this bursting behavior is explained by fluctuations in secondary structure, we confirmed that the bursting is still evident in the presence of a blocking oligo that hybridizes to the 3’NS region of miR-21 (Figure S14). Instead, we propose that the bursting behavior may result from a previously observed conformational change of miRISC between target-unbound and target-bound forms. Crystal structures in the absence of an mRNA target suggest that Ago2 primarily exposes nucleotides g2–g5^10,11,35^, with access to the remaining seed nucleotides g5–g8 blocked by α-helix-7 in a conformation dubbed α7.^10^ By contrast, target-bound crystal structures show an alternate α7′ conformation that exposes g2–g8, thus permitting stable seed binding.^35^ This has been interpreted as evidence that target pairing to nucleotides g2–g5 induces helix-7 to occupy conformation α7′. However, the bursting behavior we observe is consistent with miRISC spontaneously alternating, on a timescale of tens of seconds, between conformations with either unstable or stable seed binding. In light of this, the previously observed differences between target-free (α7) and target-bound (α7′) forms^35^ may result from a protein-intrinsic conformational equilibrium, with conformational capture of α7′ upon target binding to the seed. These intrinsic dynamics may contribute to target selectivity through acceleration of seed dissociation, a role previously attributed to helix-7 through structural and mutation studies^35^. These dynamics may also facilitate post-cleavage target release by introducing strain in the system, as well as prevent cleaved product rebinding (which can otherwise happen at a diffusion limited rate constant) for extended periods to render enzyme turnover more efficient.^25^

We also identified much larger and more variable ISIs for t9_7S_ compared to t7_6S_ (Figure 3G-H). This may be related to previous observations that pairing to g9 (not conserved in miRNAs) can actually reduce target affinity, perhaps by necessitating additional conformational changes that open the central cleft.^10^ In contrast, for targets with pairing in the central region (g10–g12), the ISIs become shorter and more narrowly distributed again, perhaps because pairing in the central region weakens guide-target interactions at g9. ISIs become somewhat longer and more broadly distributed again with pairing up to g14 (t14_7s_), as pairing extending into the supplementary chamber can even start detaching the 3’end from the PAZ domain.^26^ In contrast, our analysis showed no evidence of bursting behavior for targets longer than t14_7s_. One possible explanation is the very slow dissociation rate constants of those targets, which is likely slower than the rate constant of the conformational transition. For reliable burst analysis, the fluorescent probe should have binding and dissociation rate constants that are (at least) an order of magnitude faster than the isomerization rate constant of the RNA transcript^39^, which is not the case for targets longer than t14_7s_.

### Conformational flexibility of miRISC controls seed-less 3′-non-seed interaction with target

The major parameter thought to explain the selection of the target mRNA is the base pairing between the 5′S region of the miRNA and its target mRNA. The existing understanding of miRISC targeting is that pairing with the 5′S region allows the supplementary and 3’NS nucleotides of the guide strand to interact with the target.^10,11,13^ In fact, in a 5’S-paired ternary complex, the Watson-Crick faces of the supplementary bases (g13–g16) are splayed out towards the solvent, in a manner similar to nucleotides g2–g5 of the seed region in a guide-only structure, suggesting that 5’S binding exposes the supplementary bases for fast target interrogation.^10^ However, a new class of miRNA recognition elements (MREs) have been suggested to exclusively function at the protein coding sequence (CDS) and depend on extensive 3’NS interaction without the involvement of 5’S^15–18^. Moreover, prior crystal structures indicate a mobile 3′-end of the guide miRNA that can be detached from the PAZ domain by extensive base pairing at the 5′S and supplementary region^10,36,37^. Our SiMKEPS assays provide us a means to investigate the impact of the mechanistic parameters—such as the number of base pairs formed— underlying such 3’NS interactions with the CDS. To this end, we used Cy5-labeled target mimics of variable length (7–10 nt) complementary to the supplementary (g13–g16) and the remaining 3′NS (g17– g22) regions of miR-21 but omitting any 5′S pairing (Figure 4A). We see clear evidence of repeated binding of these probes to miR-21 RISC, comprising two populations with different kinetics (transient and long-lived), as shown by the two distinct clusters in scatter plots of the median τ_bound_ and τ_unbound_ dwell times (Figure 4B). The specific interaction of these probes with Ago2-bound miR-21 (Figure 4B and 4C) differs from any nonspecific interaction with other matrix components (Figure S15) or with miR-21-free Ago2 (Figure S16).

**Figure 4.**
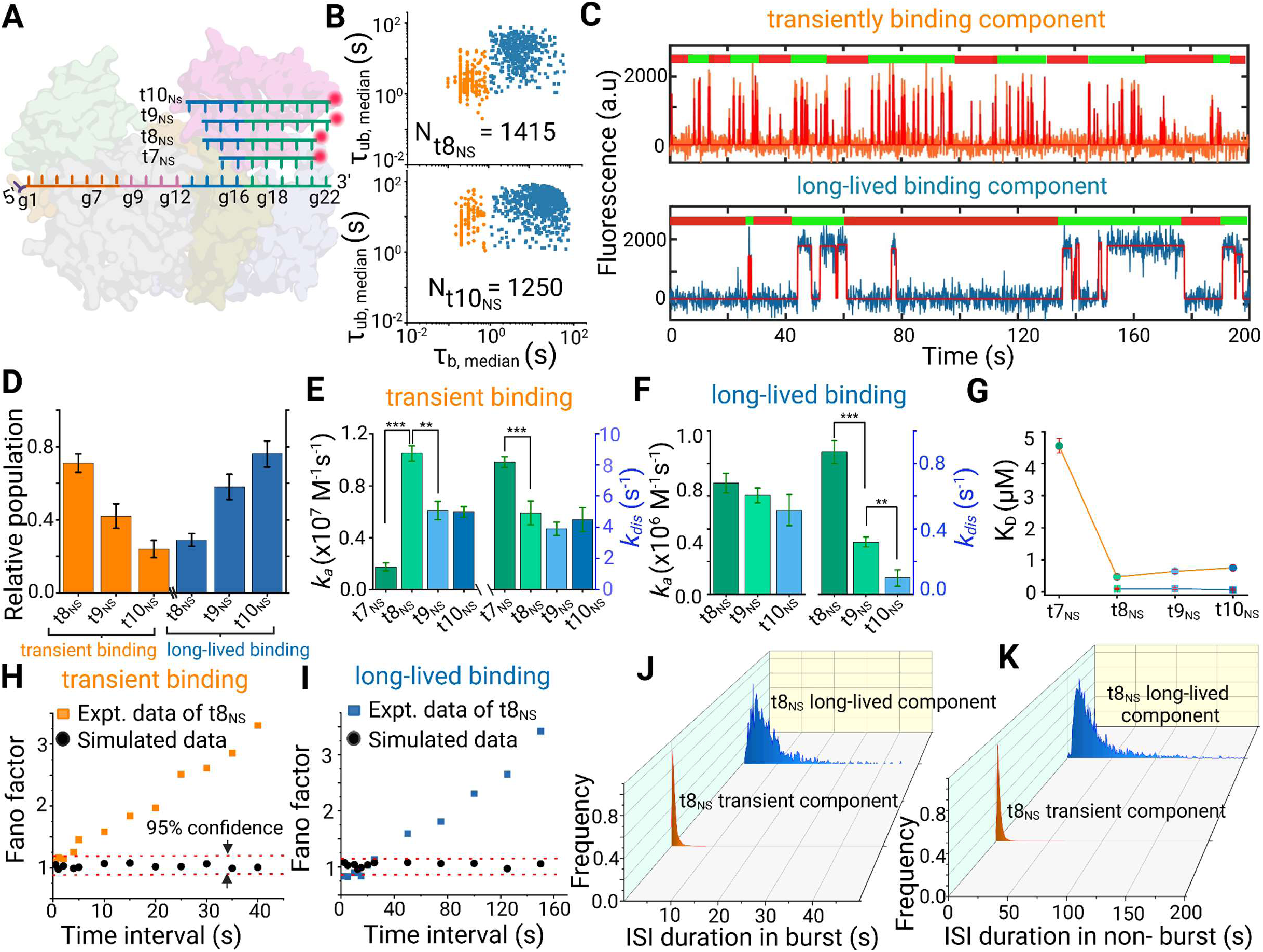
Seed-independent binding of non-seed target mimic probes to miR-21 RISC with heterogeneous kinetics (A) of the median τ_bound_ and τ_unbound_ dwell times for all intensity-versus-time trajectories within a single field of view in the presence of Cy5-labeled 8-nucleotide (t8_NS_, upper scatter plot) and 10-nucleotide (t10_NS_, lower scatter plot) target mimics interacting with miR-21 RISC. Two distinct clusters of traces characterized by different binding kinetics (blue squares and orange filled circles) are seen in each scatter plot with respective total accepted molecules (N) indicated. (**C**) Two representative intensity-versus-time trajectories showing distinct kinetics of interaction of the same non-seed probe (t8_NS_) with different copies of miR-21 RISC. The upper trace (orange) shows transient binding events, while the lower trace (blue) shows long-lived binding interactions. (**D**) Normalized relative populations of the long-lived and transient binding components for the respective probes employed; mean of independent triplicates ±SEM. (**E**) Interaction kinetics of Cy5-labeled non-seed probes of varying length to miR-21 RISC. Association (*k_a_*) and dissociation (*k_dis_*) rate constants for the transiently binding component were calculated from exponential fitting to the cumulative distributions of τ_unbound_ and τ_bound_ dwell times, respectively (error bars represent the SEM from three independent replicates; Statistical significance of differences in the rate constants was determined using two-tailed Student’s test (t test). P values < 0.1 were considered significant ***p < 0.01, **p < 0.05, *p < 0.1). (**F**) Association (*k_a_*) and dissociation (*k_dis_*) rate constants for the long-lived binding component were calculated from exponential fitting to the cumulative distributions of τ_unbound_ and τ_bound_ dwell times, respectively (error bars represent the SEM from three independent replicates; ***p < 0.01, **p < 0.05, *p < 0.1). (**G**) Binding affinity (dissociation equilibrium constant, K_D_) of target mimics having varying degrees of guide complementarity to the supplementary and 3’NS region of miR-21 RISC. Mean of independent triplicates ± SEM. (**H**) Calculated Fano factor for transiently binding component across various time intervals for the interaction of miR-21 RISC and t8_NS_ target mimic indicates non-Poissonian statistics. The black dots represent the Fano factor of the simulated data. The dashed lines indicate the 95% confidence interval for Poisson behavior. (**I**) Calculated Fano factor for the long-lived binding component across various time intervals for the interaction of miR-21 RISC and t8_NS_ target mimic indicates non-Poissonian statistics. The black dots represent the Fano factor of the simulated data. The dashed lines indicate the 95% confidence interval for Poisson behavior **(J)** Cumulative histogram displaying the distribution of inter spike intervals (ISIs) during burst period for transient (orange shade) and long-lived component (blue shade) (**K**) Cumulative histogram displaying the distribution of inter spike intervals (ISIs) during non-burst period for transient (orange shade) and long-lived component (blue shade) of t8_NS_ probe interacting with miR-21 RISC.

In the case of the 8-nt probe t8_NS_, the transient binding component (Figure 4B, orange clusters) shows fast rate constants of both association (1.05 x 10^7^ M^-1^s^-1^) and dissociation (4.93 s^-1^) (Figure 4E and Table S3). This association rate constant is ∼16 times faster than to miR-21 alone (6.50 x 10^5^ M^-1^s^-1^), suggesting that, like the seed region of miR-21 RISC, the supplementary and 3′NS nucleotides are pre-organized by Ago2 to bind to an RNA target (Figure 4E, S18). Reducing the length of the probe to 7 nt (t7_NS_(1), t7_NS_(2), or t7_NS_(3)) yields a nearly five-fold drop in association rate constant (∼0.2 x 10^7^ M^-1^s^-1^), irrespective of the sequence (Figure 4E, Figure S17 and Table S3), indicating that a contiguous stretch of ≥8 base pairs is necessary for efficient docking to the non-seed part of miR-21 (Figure 4E and Figure S12 and Table S2). The association rate constant does not increase further when the target length is extended to 9 nt (t9_NS_) or 10 nucleotides (t10_NS_). In fact, we observe a slight (∼2-fold) decrease in the association rate constant for t10_NS_ (0.60 x 10^6^ M^-1^s^-1^) compared to t8_NS_ (Figure 4E). In the case of the long-lived binding component (Figure 4B, blue clusters), the association rate constant (∼7.0 x 10^5^ M^-1^s^-1^) is ∼10-15 times slower than that of the transiently binding component (Figure 4F and Table S3 and S4), and comparable to the association rate constant to miR-21 alone (6.50 x 10^5^ M^-1^s^-1^; Figure S18). This is consistent with the long-lived binding component representing a population in which Ago2 does not pre-organize the 3′-end of the guide strand for target binding, which we hypothesize is due to detachment of the 3′-end from the PAZ domain.

If the transiently binding and long-lived components represent two miRISC populations distinguished by the extent of interaction between the PAZ domain and the 3′-end of the guide RNA, their dissociation kinetics from the target should also exhibit markedly different dependence on the number of base pairs. For the transiently binding component, the dissociation rate constant is nearly invariant over the range of 8-10 base pairs with the target (Figure 4E). This is contrary to expectations from simple hybridization thermodynamics, which predict slower dissociation as the number of base pairs is increased, and is again consistent with Ago2 chaperoning interactions with the 3′-end of the guide RNA. Moreover, the very short binding interactions for the transient population—130 times shorter than those of the same probe (t8_NS_) to a bare miR-21 (Figure 4E and Figure S18)—indicates that Ago2 perturbs the interaction between miRNA and target such that not all the nucleotides of the guide strand’s 3′-end can stably pair to the target, perhaps due to sequestration of the 3′-end by the PAZ domain.^10^ In contrast, the dissociation rate constant of the long-lived component decreases as the target length increases from 8 to 10 nt (Figure 4F), as expected for simple RNA-RNA hybridization and thus consistent with the 3′-end detaching from the PAZ domain.

If the long-lived binding component represents a population of miRISC in which the guide RNA is detached from the PAZ domain, targets with greater complementarity with the 3′-end should result in more frequent detachment and, thus, a higher population of the long-lived component. Indeed, increasing the non-seed target length from 8 to 10 nt results in an increasing fraction of miRISC with long-lived binding (Figure 4D). For example, the 8-nt probe t8_NS_ (complementary to guide nucleotides g14–g21) shows a ∼3-fold higher fraction of transiently binding complexes than the 10-nt probe t10_NS_ (complementary to guide nucleotides g13–g22) and a correspondingly lower fraction of long-lived binding (Figure 4D). This is consistent with the prior finding that just one additional base pair with the target mRNA detaches the 3′-end of the guide strand from the PAZ domain, as is the case with bacterial TtAgo.^40,41^

Based on our observation of bursting behavior in 5′S binding, as well as the known coupling between helix-7 (influencing seed accessibility) and the PAZ domain (influencing 3′NS accessibility), we also expected bursting behavior in 3′NS binding as a result of intrinsic conformational fluctuations of Ago2. Indeed, burst analysis reveals non-Poissonian statistics of 3′NS probe-binding dwell times and suggests that a slow (∼tens of seconds) equilibrium conformational transition periodically permits binding of non-seed probes (Figure 4H–4K). The similarity in bursting behavior between the 5′S and 3′NS probes suggests that the same (or similar) conformational dynamics of Ago2 might influence accessibility of both the seed and non-seed/supplementary regions of the guide RNA.

### miRISC exhibits multiple stable states with mutually exclusive 5′S and 3′NS accessibility

We hypothesized that the bursting behavior we see for both 5′S and 3′NS binding is linked to conformational changes of helix-7 and the PAZ domain, which are known from prior work to move as a rigid body within Ago2.^10–13^ This hypothesis predicts that the binding of each miRISC to 5′S and 3′NS target mimics will not be independent, but correlated. To test this prediction, we performed two-color measurements to simultaneously observe the binding of both a 10-nt Cy3-labeled seed probe (t10_7S_) and an 8-nt Cy5-labeled non-seed probe (t8_NS_) to miR-21 RISC (Figure 5A). These measurements unveiled three distinct clusters in the scatter plot of the median τ*_bound_* and τ*_unbound_* dwell times (Figure 5B). One cluster (in green) corresponds to the binding of the Cy3-labelled t10_7S_ probe to the 5’S region of miR-21 RISC. The other two clusters (orange and blue clusters) represent the transient and long-lived binding, respectively, of the Cy5-labelled t8_NS_ probe to the supplementary plus 3’ NS region. Of all accepted traces in both the Cy3 and Cy5 channels, ∼18% of traces show transient binding of t8_NS_, 30% show long-lived supplementary plus 3’NS binding kinetics of t8_NS_ and 72 % show kinetics akin to the 5’S binding of t10_7S_ (Figure 5B and 5C).

**Figure 5.**
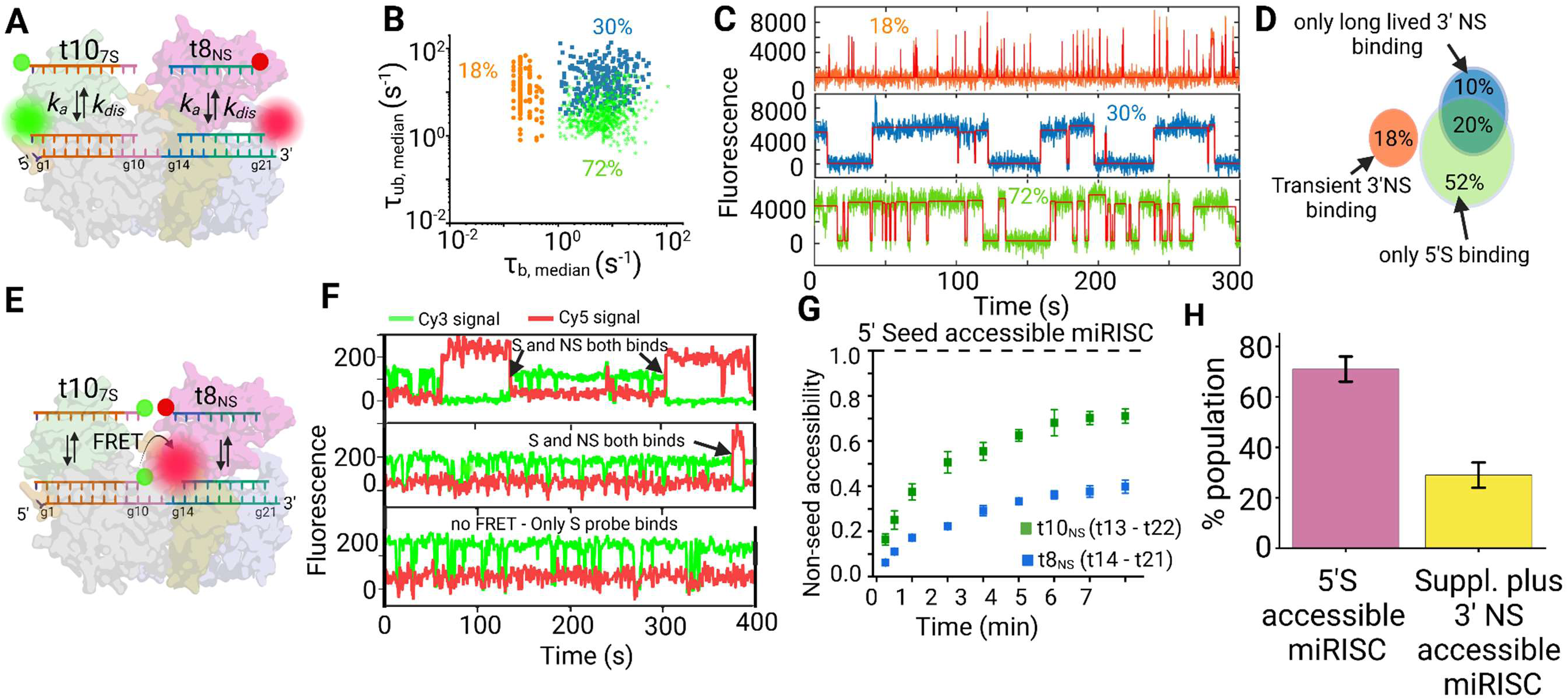
Seed and non-seed binding within single miRISC complexes. (**A**) Schematic showing the design of Cy3-labeled seed (t10_7S_) and Cy5 labelled non-seed probes (t8_NS_) employed to simultaneously investigate accessibility of seed and non-seed regions of miR-21 RISC. (**B**) Scatter plot of the median τ_bound_ and τ_unbound_ dwell times when both seed (t10_NS_) and non-seed probe (t8_NS_) are employed simultaneously. The scatter plot shows three distinct clusters whose kinetics are consistent with miRISC populations accessible to either the seed (seed cluster) or non-seed (orange and blue clusters) probe. (**C**) Representative intensity *versus* time traces of miR-21 RISCs whose chromato-kinetic properties are suggestive of interaction with t8_NS_ (upper and middle traces), and t10_7S_ (lower trace) within the observation time window of 300 s. The relative percentage of each kind of traces accepted in the collected movie are mentioned (**D**) Venn diagram showing that a 20% of long-lived non-seed binding component has also seed accessibility and vice-versa. However, the transient non-seed component and a fraction of long-lived non-seed binding components has no seed accessibility. **(E)** Schematic showing the design of FRET probes for probing simultaneous seed and non-seed accessibility. The 10-nt seed probe is labeled with Cy3 to serve as a FRET donor, and a Cy5-labeled 8-nt non-seed probe acts as a FRET acceptor. (**F**) Representative FRET trajectories showing simultaneous seed and long-lived supplementary plus non-seed binding evident from anti-correlated changes in Cy3 and Cy5 signal. (**G**) Plot showing the fraction of seed-accessible miR-21 RISCs that also show binding to the supplementary and non-seed region by the t8_NS_ or t10_NS_ probe within varying observation periods up to 7 min in length. Error bars represent the SEM from two independent replicates **(H)** The percentage of different miRISC conformations present in cell extract. Reddish purple bar represents the percentage of miRISC (∼70%) that show stable 5’S binding and yellow bar show the percentage of supplementary and 3’NS binding miRISC population (∼30%, sum of transient and long-lived binding component) that do not show 5’S binding in our experiment.

Colocalization analysis of fluorescent spots in the Cy3 and Cy5 channel showed that the vast majority of traces with transient 3′NS binding (∼97%) do not exhibit any 5’S binding, suggesting that these two populations represent states of Ago2 with mutually exclusive seed and non-seed binding (Figure 5B, 5C and Figure S19), consistent with our hypothesis that coupled conformational changes of helix-7 and PAZ domain result in correlated changes in seed-and non-seed accessibility. However, a considerable fraction (∼ 20%) of the miRISC population displays both 5’S and long-lived 3’ NS binding (Figure 5D).

To more directly confirm that seed and non-seed probes can simultaneously bind the same miRISC complex, we next performed FRET experiments with a 5′-Cy3-labeled 10-nt (t10_S_) seed probe as the FRET donor and 3′-Cy5-labeled 8-nt or 10-nt non-seed probe as the acceptor, chosen to position the two fluorophores in close proximity with one another when both probes are bound (Figure 5E). The 10-nt-long Cy3-labeled donor seed probe was chosen due to its long bound-state dwell times, which increases the probability of FRET to the more transiently binding Cy5-labeled acceptor probe. Occasional dissociation and re-binding replenishes the donor Cy3 probe with fresh probe from solution, reducing the influence of dye photobleaching and allowing for the observation of probe-binding events over an extended period of several minutes. When using a 10-nt non-seed probe (t10_NS_), 99% of the observed FRET traces display long-lived non-seed binding kinetics (Figure 5F), consistent with our previous observation that only the long-lived 3′NS binding population shows seed binding. Of those miRISCs exhibiting 5′S binding, approximately 70% also exhibit long-lived non-seed probe binding with the remaining 30% showing no evidence of binding in the supplementary and 3′NS region within the 7-minute observation period (Figure 5G and Figure S20). However, with the 8-nt non-seed probe (t8_NS_), only 40-45% of traces exhibit any FRET transitions corresponding to non-seed binding (Figure 5G). The remaining 55–60% of traces show seed probe binding but no FRET and, hence, no evidence of non-seed probe binding (Figure 5G). The higher yield of long-lived non-seed binding for t10_NS_ than t8_NS_ aligns with the hypothesis (based on observations in Fig. 4D) that a 10-nt non-seed probe, fully complementary to the supplementary and the 3′ non-seed regions, is more efficient in displacing the 3′-end from the PAZ domain than an eight-nucleotide non-seed probe. In this interpretation, we only see FRET signal from simultaneous binding of seed and non-seed probes when the 3′-end of the guide strand disengages from the PAZ domain of a 5′S-accessible miRISC. Notably, the association rate constant of the t8_NS_ probe to the supplementary and 3′NS region (0.30 x 10^6^ M^-1^s^-1^) is 14-fold slower than the binding of the 10-nt seed probe (t10_7S_) to the 5′S region (4.2 x 10^6^ M^-1^s^-1^) (Table S2 and Table S4), which aligns with the conventional view that the seed interaction is established before proceeding to extended base pairing in the supplementary and 3’NS regions.

Thus, the combined results of SiMKEPS and smFRET analysis, while generally consistent with the standard model of miRNA target recognition, also provide evidence of subpopulations of miRISC with exclusive 5′S or 3′NS accessibility that may be the result of intrinsic conformational dynamics of Ago2, in particular the correlated movement of helix-7 and the PAZ domain. We conclude that a considerable fraction of cellular miRISC (28 ± 5%) (sum of transient and long-lived component) exists in an additional stable conformational state with supplementary and 3′NS accessibility but not 5′S accessibility (Figure 5H). We also show that a subset of miRISC complexes can simultaneously bind targets via the 5′S and 3′NS end, with the probability of such simultaneous binding increased with longer stretches of 3′NS base pairs (Figure 5D and 5G)

## Discussion

Cellular miRISC binds to target messenger RNA (mRNA) sequences and prevents their translation into proteins, thereby regulating gene expression. This has conventionally been understood to initiate *via* 5′S binding, which involves the binding of the seed region (g2–g8) at the 5′-end of the miRNA to the target sequence.^10,11,13^ According to the established stepwise mechanism of miRISC targeting, 5′S binding triggers a conformational change of Ago2 (involving movement of helix-7 and the PAZ domain as a rigid body) that induces further base pairing to the supplementary and 3′NS regions^10,13^. Here, using SiMKEPS we have shown that a subpopulation of miRISC exhibits transient binding to the supplementary and 3′NS region independent of—and, indeed, mutually exclusive with—seed binding. This suggests the presence of an additional stable conformation of miRISC in cells, with helix-7 and PAZ domain displaced from their typical orientation that favors seed binding into one that only allows supplementary plus 3’NS interaction with the target.

We have also shown that authentic miRISC molecules isolated from a mammalian cell extract undergo slow, spontaneous state transitions that raise or lower the accessibility of the 5′-and 3′-ends of the guide RNA. We speculate that these states arise from conformational fluctuations involving helix-7 and the PAZ domain of Ago2 which, through coupled movements, affect access of the target to both the seed region and the supplementary and 3′NS regions of the guide strand.^10^ The fact that these states persist over multiple target-mimic binding cycles raises the possibility that previously observed differences between target-bound and target-free conformations of Ago2 may arise in part from conformational capture of intrinsic Ago2 fluctuations, instead of (or in addition to) conformational changes induced by seed binding. These spontaneous excursions into less-accessible states may provide a mechanism for Ago2 to avoid overly stable binding or to enhance turnover, a role that has previously been ascribed to helix-7.^10^ We also observe a separate population of miRISC that exhibits stable interaction with the supplementary and 3′NS region that is independent of, and *not* mutually exclusive with, 5′S binding. This lack of mutual exclusivity and kinetics that strongly resemble interaction with a free miRNA are consistent with this population representing detachment of the 3′-end of the guide strand from the PAZ domain, which is promoted by longer probes that exhibit more extensive base pairing to the supplementary and 3′NS regions. Our study thus opens a fresh perspective on miRNA targeting by recognizing that, irrespective of 5′S binding, the supplementary and 3′NS region of the miRNA can engage in at least two independent modes of RNA sequence recognition.

These results lead us to propose a unifying model of miRISC targeting (Figure 6) wherein the miRISC can identify its target mRNAs via two distinct mechanisms: 5′S and supplementary plus 3′NS-based target recognition. These two mechanisms are independent of each other and are mediated by different stable conformational states of miRISC. Previous reports have indicated that the majority of miRISC populations exist in a conformation wherein the 5′S region is accessible for target binding, and that optimal stability of target binding is achieved through base pairing in the 5’S (g2–g8) and supplementary (g13–g16) regions, while binding to the central (g9–g13) and 3′ NS (g17–g19) regions does not correlate with the affinity of target binding.^10, 28^ Consistent with these previous reports, for the majority of miR-21 RISCs with an accessible 5′S (∼75%) we find that binding of seed-only probes is transient, but can be stabilized by additional base pairing in the supplementary region (g13–g16). However, the supplementary interaction provides only a modest increase in affinity as base pairing to the supplementary region also reduces the target association rate constant due to the possible detachment of 3’-end from PAZ domain. In addition, the slower dissociation upon supplementary base pairing may reflect an avidity effect of two otherwise largely independent base pairing interactions.

**Figure 6:**
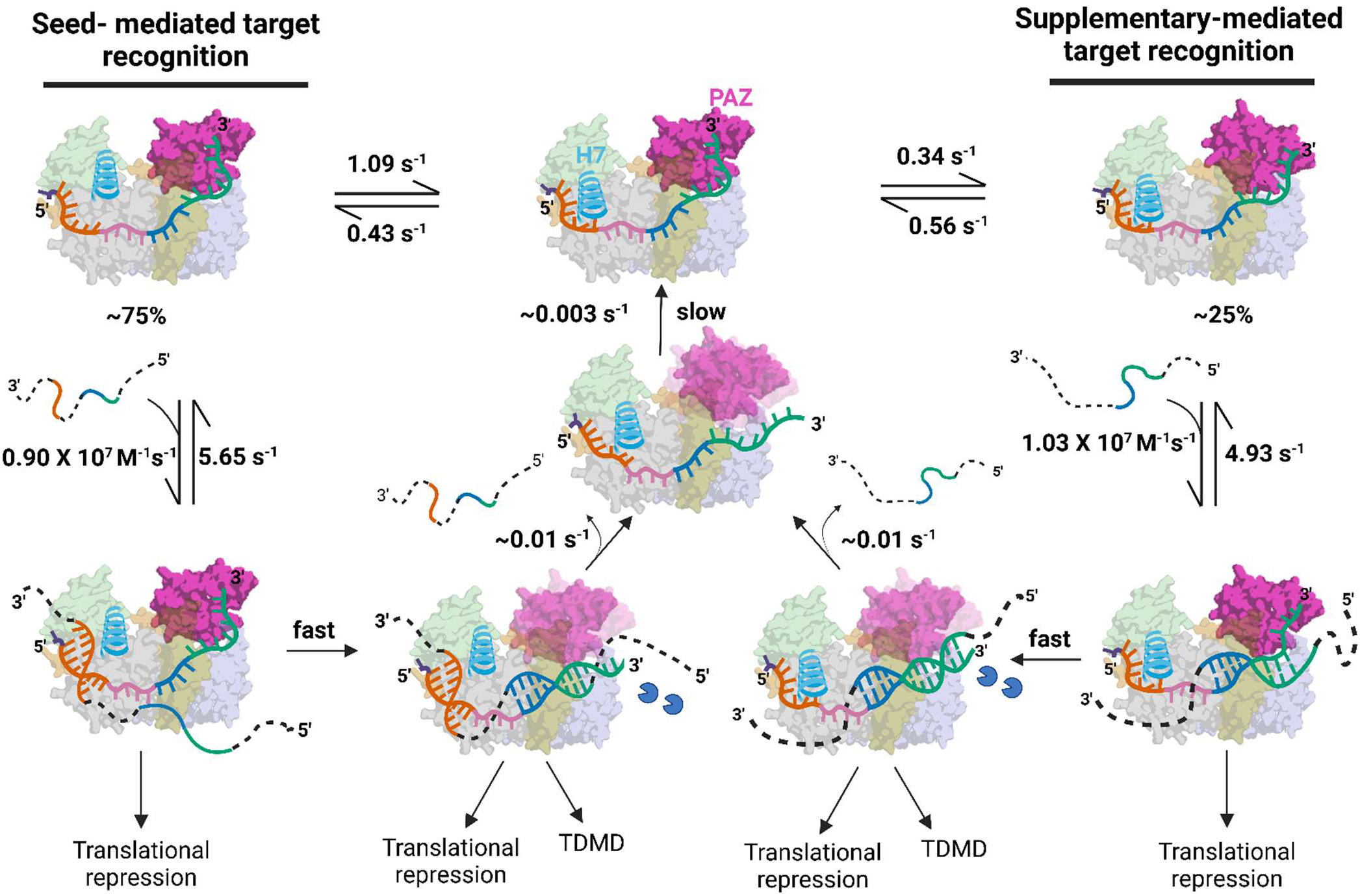
Resulting unifying model of miRISC targeting of mRNA.

miRISCs showing only supplementary and 3′-NS accessibility (∼25%) can be assigned to a conformation in which the 3′NS end is still attached to the PAZ domain but can engage in conformational fluctuation-dependent transient non-seed interactions with the target mRNA (Figure 6). In this model, since the 5′S region is inaccessible for stable target binding due to the unfavorable orientation of α-helix-7^35^ and the 3′NS end is still associated with the PAZ domain, the supplementary chamber (g13-g16) could serve as the seeding region for target interrogation in this miRISC conformation. However, extensive base pairing to the supplementary and 3′NS region can displace the 3′-end from the PAZ domain, resulting in completely different kinetics of target interaction (Figure 6). This detachment of PAZ, previously reported to be induced by pairing with g17–g21, can lead to formation of a TDMD complex whose 3′-end is stably displayed for enzymatic attack.^37,42^

The existence of target sites that reside in the target protein-coding sequence (CDS) and lack seed complementarity but have extended pairing to the 3′NS-end of miRNAs has been reported and functionally validated.^15–18^ Our results show that, compared to 5′S interactions, target interrogation through the 3′NS region is either transient or, if long-lived, possesses a slow association rate constant (Figure 6). This, together with fewer 3′NS accessible miRISCs, may explain why targeting through seed-less 3′NS interactions results in only a modest effect on mRNA stability and protein translation.^15–18^ Nevertheless, our results show that such binding can occur, and may still yield significant cumulative effects on miRNA regulation of mRNA expression in cells. In fact, CLASH found that miRNAs that interact with targets through the supplementary and 3′NS regions tend to have higher GC content^15^, which may in part compensate for the lower efficiency we and others observe for this mode of targeting.

In summary, our results illustrate the surprisingly multifaceted modes of miRISC-target interactions, revealing a detailed picture of how the 3′NS region can interact with target mRNA sequences independently of 5’S binding. Many miRNAs exhibit especially high sequence conservation within the seed region (g2–g8) and nucleotides g13–g19 of the supplementary and 3′NS regions.^15^ We speculate that each of these conserved regions may, *via* distinct miRISC conformations, independently or additively contribute to regulation of mRNA expression by targeting distinct regions of mRNAs (*i.e.*, 3′ UTRs or CDS). The unifying model of miRISC targeting of mRNA proposed here reconciles previously divergent findings and sheds light on a previously unknown mechanism of miRISC function relevant to RNAi therapies.

### Limitations of the study

Although this study identified a unique conformation of miRISC that allows for supplementary and 3’NS interactions with targets, the precise arrangement of the protein subunits, particularly the exact orientation of the α-helix-7 and PAZ domains, remains to be elucidated in future structural investigation. Additionally, whether chemical modifications of either the miRNA or Ago2 within cells are required to favor this unique miRISC conformation is also a subject for future study. We attempted to prepare miR-21 RISC complexes using purified RNA-free AGO2 and hsa-miR-21-5p, but obtaining these recombinant complexes proved challenging, as seems typical for the field. Nevertheless, the formation of these complexes of unique conformation in structural studies suggests that they can form spontaneously in solution without the need for other cellular components.

## STAR ★ METHODS

### Detailed methods are provided in the online version of this paper and include the following

- KEY RESOURCES TABLE
- **EXPERIMENTAL MODEL AND STUDY PARTICIPANT DETAILS**

ᐤ HeLa cell
- METHOD DETAILS

ᐤ Biotinylation of capture antibody
ᐤ Fluorophore labeling of detection antibody
ᐤ Preparation of slide surfaces for single-molecule microscopy
ᐤ Total Internal Reflection Fluorescence (TIRF) microscopy
ᐤ Preparation of imaging solution
ᐤ Kinetic fingerprinting assays of miRISC (SiMKEPS)
ᐤ Analysis of kinetic fingerprinting data
ᐤ Fitting of the cumulative frequency of dwell times
ᐤ Förster Resonance Energy Transfer (FRET) experiment and data analysis
ᐤ Fano factor calculation
ᐤ Burst analysis
- QUANTIFICATION AND STATISTICAL ANALYSIS

## Acknowledgements

This work was funded by NIH grant GM131922 to N.G.W. SiMREPS analysis software for kinetic fingerprinting analysis was developed in part based on funding from a Michigan Economic Development Corporation MTRAC for Life Sciences grant to N.G.W. and A.J-B, as well as NIH grants R21 CA204560 and R33 CA229023 to N.G.W.

## Author contributions

T.C, S.M, and N.G.W. conceived the idea and devised the approach for kinetic fingerprinting-based accessibility detection of guide strand of miRISC. T.C., S.M. and S.R. designed the experiments, collected, and analyzed the data, and wrote the initial draft of the manuscript. A.J.B. performed the trace simulations and single-trace kinetic analyses. All authors interpreted the data and edited the manuscript.

## Additional information

Supplementary information is available in the online version of the paper. Reprints and permissions information is available online at www.cell.com. Publisher’s note: Cell remains neutral with regard to jurisdictional claims in published maps and institutional affiliations. Correspondence and requests for materials should be addressed to N.G.W.

## Data Availability

All data supporting the findings of this study are available within the paper and its Supplementary Information. Source data as well as all MATLAB script are available through DeepBlue deposit.

## CORRESPONDING AUTHOR

Correspondence to Nils G. Walter.

## STAR ★ METHODS

### KEY RESOURCES TABLE

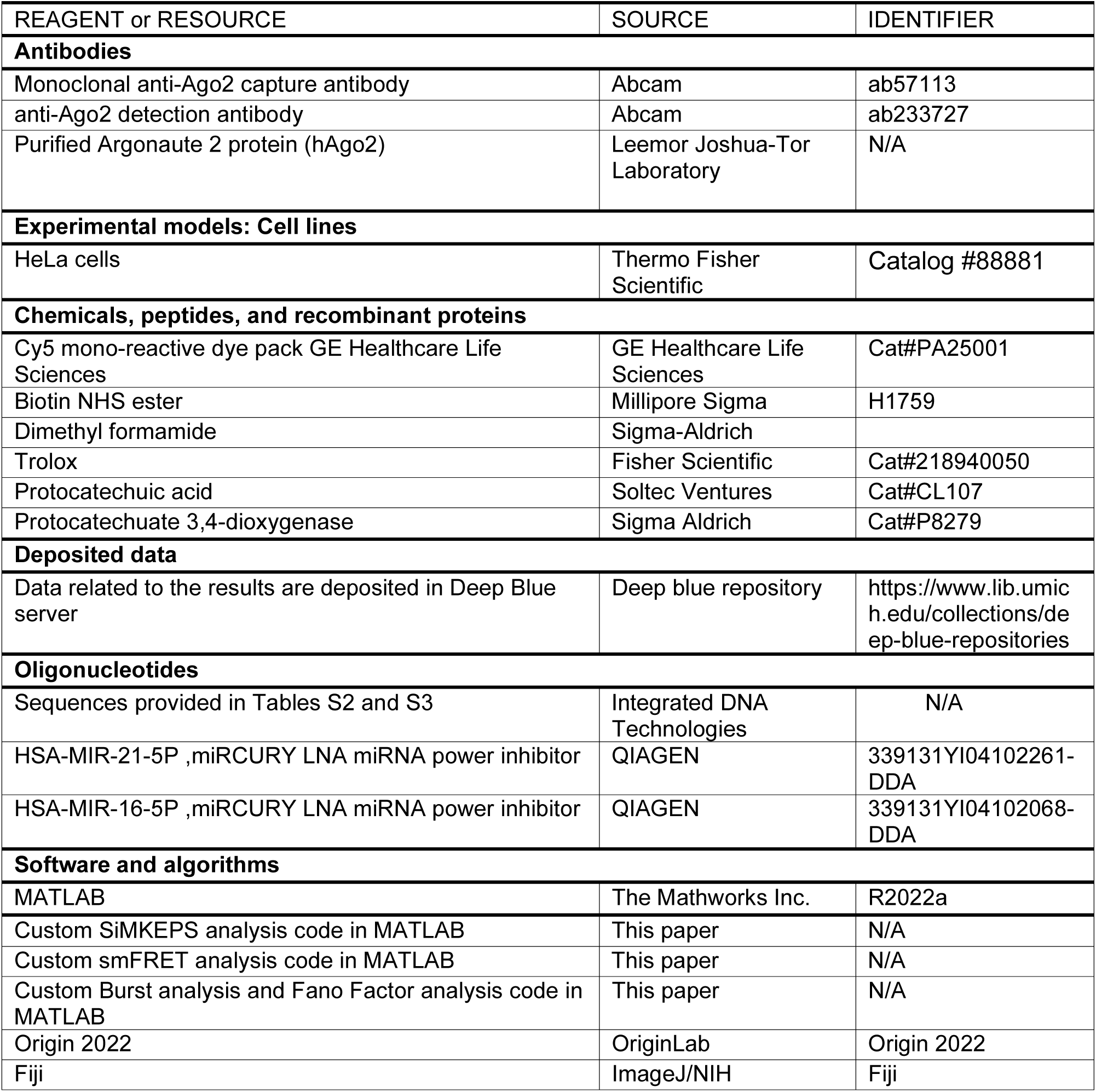

## EXPERIMENTAL MODEL AND STUDY PARTICIPANT DETAILS

### Cell extract preparation

HeLa whole cell lysate was purchased from Thermo Scientific. These cells were then centrifuged at 12000 rpm to remove the cell debris. the clear supernatant was then transferred to another Eppendorf tube before transferring it to the sample chamber.

## METHOD DETAILS

### Biotinylation of capture antibody

Monoclonal capture antibodies were biotinylated by amine-NHS ester coupling using biotin *N*-hydroxysuccinimidyl ester (Sigma Aldrich, Catalog #H1759-100) in reactions containing a biotin:antibody molar ratio of 5:1 which were carried out at room temperature for 1 h in 1× PBS, pH 7.4. Biotin-IgG conjugates were purified using Zeba Spin desalting columns (ThermoFisher, Catalog #89882, 7K MWCO) according to the manufacturer’s recommended protocol, followed by overnight dialysis at 4 °C (Slide-A-Lyzer Dialysis Cassette, ThermoFisher, 3.5K MWCO) against 1× PBS, pH 7.4. The fraction of biotinylated IgG was estimated by electrophoretic mobility shift assay in the presence or absence of excess streptavidin and ranged from 70% to 80%. Capture antibodies were aliquoted and frozen at-80°C.

### Fluorophore labeling of detection antibody

Candidate detection antibodies were fluorescently labeled by amine-NHS ester coupling using Cy5 mono-reactive dye packs (GE Healthcare, Catalog #PA25001). A molar dye:protein ratio of 10:1 was used in labeling reactions, which were carried out in the dark at room temperature for 1 h. Antibody-dye conjugates were purified using Zeba Spin desalting columns (ThermoFisher, Catalog #89882, 7K MWCO) according to the manufacturer’s recommended protocol, followed by overnight dialysis at 4 °C (Slide-A-Lyzer Dialysis Cassette, ThermoFisher, 3.5K MWCO) against 1× PBS, pH 7.4. The ratio of dye:antibody after purification was quantified by UV-Vis spectrophotometry (Nanodrop) using the absorbance values at 280 nm and 650 nm and was measured to be 3:1.

### Preparation of slide surfaces for single-molecule microscopy

Glass coverslips (No. 1.5, 24 x 50 mm, VWR Catalog #48393-241) were functionalized with a 1:100 mixture of biotin-PEG-SVA and mPEG-SVA (Laysan Bio, Inc., Catalog #MPEG-SVA-5000-1g and #BIO-PEG-SVA-5K-100MG) as previously described.^43^ The coverslips are passivated with methoxy polyethylene glycol (mPEG) to minimize nonspecific binding as well as biotinylated PEG which, *via* streptavidin, binds the biotinylated anti-Ago2 antibody. Coverslips were stored under aluminum foil in a nitrogen-purged cabinet until use (up to 4 weeks). Prior to an experiment, 2-6 sample cells were attached to each coverslip by cutting a ∼2-cm length from the wider end of micropipette tips (Thermo Fisher, #02-682-261), discarding the narrower segment of the pipet tip, and then placing the wide end down on the PEGylated glass coverslip and then sealing the edges with epoxy adhesive (Ellsworth Adhesives, #4001).

### TIRF microscopy

SiMREPS experiments were performed using Olympus IX-81 objective-type TIRF microscopes equipped with cellTIRF and z-drift control modules (ASI CRISP) and Oxford Nanoimager (ONI). Cy5-labeled oligonucleotides and antibodies were excited in TIRF mode with a theoretical penetration depth of ∼80 nm using a fiber-coupled diode laser (OBIS 637nm LX, 100mW) with an incident light intensity of ∼5 W/cm^2^, and fluorescence emission was detected using an EMCCD (Photometrics Evolve) with an exposure time of 20 to 250 ms, after passing through a dichroic mirror and emission filter (ET655LP-TRF). In some experiments, an objective heater (Bioptechs) was used to raise the observation temperature (calibrated against a reference thermistor provided by the manufacturer for the specific sample cell geometry used in this study).

### Preparation of imaging solution

Unless otherwise specified, all kinetic fingerprinting and FRET assays were carried out in an imaging solution comprising 1× PBS, pH 7.4 (Gibco); an oxygen scavenger system^43^ consisting of 5 mM 3,4-dihydroxybenzoic acid (Fisher, #AC114891000), 0.05 mg/mL protocatechuate 3,4-dioxygenase (Sigma Aldrich, #P8279-25UN), and 1 mM Trolox (Fisher, #218940050); and 50 nM fluorophore-labeled oligonucleotide. In experiments with multiple probes (2 × Cy5 or Cy3 + Cy5), each probe was at 50 nM.

### Kinetic fingerprinting assays of miRISC

All sample handling was performed in GeneMate low-adhesion 1.7-mL microcentrifuge tubes. The slide surface was washed with 100 µL of T50 buffer (10 mM Tris-HCl, 50 mM NaCl, 1 mM EDTA, pH 8.0) for 10 minutes followed by the addition of 40 µL 1 mg/mL streptavidin. After 10 min, excess streptavidin was removed, and the sample chamber washed three times with 100 µL of 1× PBS. Next, the coverslip was coated with the biotinylated anti-Ago2 capture antibody by adding 40 µL of a solution containing 10 nM of biotinylated capture antibody in 1× PBS buffer and incubating for 30 min. Excess antibody was removed and the sample wells washed three times with 100 µL of 1× PBS. HeLa whole cell lysate was diluted 20-fold in 1× PBS buffer and then centrifuged at 9000 for 6 min at 4 degree C to remove cell debris, yielding HeLa whole cell extract. The supernatant was collected in a new microcentrifuge tube. A 100-μL portion of the cell extract in 1× PBS was added to the sample chamber and incubated for 1 h to capture the miRISC on the coverslip surface. Excess cell extract was removed from the sample well. The sample cell was then washed twice with 100 µL of 1× PBS, and 200 µL of imaging solution added. Samples were immediately imaged by TIRF microscopy.

### Analysis of kinetic fingerprinting data

Kinetic fingerprinting data were analyzed using custom MATLAB code to identify sites of fluorophore-labeled oligonucleotide probe binding and analyze the kinetics of repeated binding as described previously, using a diffraction-limited analysis pipeline^20^. Briefly, regions of repeated probe binding and dissociation (regions of interest, ROIs) in the FOV were identified by determining the average absolute frame-to-frame change in intensity at each pixel to create an intensity fluctuation map and then defining ROIs as the 3×3-pixel regions centered on local maxima within the fluctuation map. Next, the integrated, background-subtracted intensity within each ROI was calculated for each frame in the movie to generate an intensity-*versus*-time trace. These candidate traces were subjected to hidden Markov modeling (HMM) using a version of vbFRET^43^. The idealized trace generated *via* HMM was used to determine several parameters for kinetic fingerprinting analysis: N_b+d_, the number of binding and dissociation events; τ_on,median_ and τ_off,median_, the median dwell times in the probe-bound and probe-unbound states, respectively; τ_off,max_, the maximum dwell time in the probe-unbound state; and ^S^/_n_, the signal-to-noise ratio, defined as the standard deviation of the fluorescence intensity divided by the mean intensity difference between bound and unbound states. Threshold values for each of these parameters to count a trace as a positive detection event were optimized using optimizer software for each oligonucleotide probe.

### Fitting of the cumulative frequency of dwell times

Fitting an exponential cumulative distribution function (CDF) to the all-individual dwell times in the bound state (τ_b_) and unbound state (τ_ub_), obtained from Hidden Markov Modelling (HMM) of individual traces, using a single exponential function (Equation S1) or a double exponential equation (Equation S2) in Origin Lab software provides association and dissociation rate constant.

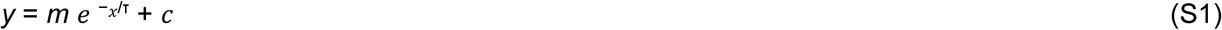

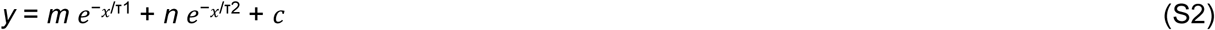

where *m, n, c*, τ, τ1 and τ2 are fit parameters for S1 and S2. The coefficients *m* and *n* are used to fit the function and for the double exponential, determine the weight of each term for plotting, and obtaining average dwell times. The coefficient τ describes, for the single exponential fit, the average dwell time for a given event. The coefficients τ1 and τ2 describe, for the double exponential fit, the average dwell time for shorter-and longer-lived populations of events, respectively. The coefficient c is a constant that gives the y-intercept for the equation. For each dataset, the cumulative frequency was first fit to the single exponential fitting function. This fit was then kept if the sum squared error < 0.05 and the R^2^ > 0.99 with a good residual fitting which indicated a good fit and suggested that the coefficient τ was an accurate average dwell time. If these conditions were not met, a double exponential function (equation S2) was used instead, and the average dwell time was calculated as τ = (*m* τ1+ *n* τ2) / (*m* + *n*). This equation calculated a weighted average of both populations that was reported as the average dwell time for the entire data set.

### Single-molecule trace simulations and single-trace fitting of dwell time distributions

Two-state SiMKEPS traces were simulated using a kinetic Monte Carlo model written in MATLAB. First, ideal noiseless traces were simulated by assuming a single average lifetime in the bound (1.4 s) and unbound (3.5 s) states, chosen to match the mean per-trace lifetimes as determined from exponential fitting of individual traces of t8_7s_ binding to miR-21 RISC (note: due to heterogeneity, these do not perfectly correspond to the rate constants in Table S2, which were determined from cumulative dwell time distributions pooled from all traces in a given condition). These lifetimes were converted from units of seconds into units of movie frames by dividing by the experimental exposure time (0.1 s), and then into per-frame transition probabilities according to the relationships

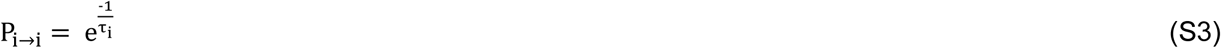

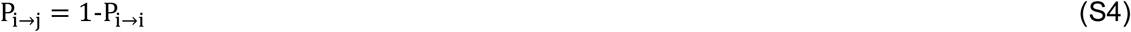

where P_i→i_ is the probability of remaining in state i in the next frame, τ_i_ is the lifetime of state i in frames, and P_i→j_ is the probability of transitioning from state i to state j in the next frame. For each frame, a pseudo-random number generator was used to choose a number n on the interval [0,1]. If n > P_i→i_, a transition to state j occurred; otherwise, the molecule remained in state i. Upon generation of the entire trace (500-8000 frames), the states (0 or 1) were multiplied by the typical intensity value of SiMKEPS traces (4000 counts) and simulated Gaussian noise with σ = 0.1 × 4000 = 400, corresponding to a typical signal-to-noise ratio of ∼10, was added to each trace. Traces were then subjected to the same HMM analysis as experimental traces to extract apparent dwell times for further analysis.

Rate constants were estimated for single traces by single-exponential fitting as follows. After extraction of individual dwell times in the bound and unbound state by HMM, and exclusion of the first and last dwell times (which are truncated), the cumulative distribution function of the dwell times in the bound and unbound state were generated for each trace and fit by a single-exponential function to extract lifetimes in the bound and unbound states. For experimental data, only those molecules passing the filtering criteria for intensity, signal-to-noise, and lifetime were included. Across the population of molecules (experimental or simulated), the coefficient of variation (CV) of each lifetime was calculated as the standard deviation divided by the mean of the lifetime.

### Förster Resonance Energy Transfer (FRET) experiment and data analysis

smFRET studies of miRNA accessibility in RISC were also performed using TIRF microscopy^44^. Emission from both Cy5 and Cy3 fluorophores was simultaneously recorded using two electron-multiplying charge-coupled device cameras (Andor iXon Ultra EMCCDs) at 250 or 50-ms time resolution using Micro-Manager software (https://www.micro-manager.org/). Fluorescence intensity *versus* time traces were extracted from the raw movie files using IDL (Research Systems) software and analyzed using MATLAB scripts.^43^

### Burst analysis

Burst analysis was carried out using the RS method described by Gourévitch and co-workers.^33^ We utilized a modified MATLAB implementation of the RS method based on the MATLAB script provided in the supplement to Gourévitch et al. In the standard implementation, each molecule’s ISIs are ranked independently. In our implementation, termed global burst analysis, we have extended this so that each ISI detected in all our experimental conditions is ranked simultaneously; therefore, a burst is defined as a global property of the molecules and is not biased by the number of binding events of an individual molecule. Briefly, ISIs were determined by calculating the time in between consecutive binding events for each molecule. ISIs for all molecules were collected and used as an input for global burst analysis, using the RS method to demarcate the start and end points of bursts—that is, the first and last binding events, respectively, in a sequential series of binding events occurring in quick succession (‘burst’). Each start and end point were then reassigned to the corresponding molecule, preserving the single-molecule burst profile. The RS algorithm developed by Gourévitch et al. requires two parameters, the maximal ISI between spikes to be considered part of a burst and an RS cutoff (alpha). These were set to 40 s and 3, respectively. Although the maximal ISI in a burst was set to 40 s, the distribution of ISIs we obtained after analysis is much lower than this, suggesting that we have provided enough flexibility in the algorithm to find the true distribution of ISIs in a burst. MATLAB scripts for global burst analysis are provided as part of the Supplementary Information.

### Fano factor calculations

MATLAB scripts were written to calculate and simulate the Fano factor from our experimental data and from a simulated Poisson distribution, respectively. The Fano factor is defined as the variance in spike counts divided by the mean spike count for a given time interval, T. For every molecule analyzed in a particular condition, a time interval of length T was randomly selected from the molecule’s fluorescence time trace. The Fano factor was calculated for T = 5, 10, 20 and 40 s. Each time window T was sampled 100 times with a different random seed for each molecule to generate an average Fano factor. For the simulations, the MATLAB Poisson random number generator was utilized to generate spike counts with an average firing rate constant equal to the average ISI in the bursts from our experimental data and an equal number of samplings. 95% Confidence intervals were calculated in MATLAB utilizing the function: bounds ¼ gaminv([.025,.975],(n 1)/2,2/(n 1)), where n ¼ sample size

## REFERENCES

1. Bartel D. P. Metazoan MicroRNAs, Cell 173, 20–51, Doi: 10.1016/j.cell.2018.03.006 (2018)

2. Friedman RC, Farh KK-H, Burge CB and Bartel D. P. Most mammalian mRNAs are conserved targets of microRNAs. Genome Res 19, 92–105. (2009)

3. Berezikov, E., and Plasterk, R.H. Camels and zebrafish, viruses and cancer: a microRNA update. Hum. Mol. Genet. 14 (Suppl 2), R183– R190. (2005)

4. Landgraf, P. et al. A mammalian microRNA expression atlas based on small RNA library sequencing. Cell 129, 1401–1414, doi:10.1016/j.cell.2007.04.040 (2007).

5. Pitchiaya, S., Mourao, M.D.A., Jalihal, A.P., Xiao, L., Jiang, X., Chinnaiyan, A.M., Schnell, S., and Walter, N. G. Dynamic recruitment of single RNAs to processing bodies depends on RNA functionality. Mol. Cell 74, 3, 521–533. (2019)

6. Peng, Y and Croce, C. M. The role of MicroRNAs in human cancer,: Signal Transduction and Targeted Therapy 1, 15004. doi:10.1038/sigtrans.2015.4 (2016)

7. Juzwik, C. A., Drake, Zhang, Y., Paradis-Isler, N., Sylvester, A., Douglas, C., Morquette, B., Moore, C. S., Fournier, A. E., miRNA dysregulationin neurodegenerative diseases; a systematic review, Progress in Neurobiology 182, 10664, DOI: 10.1016/j.pneurobio.2019.101664 (2019)

8. Chen, D., Yang, X., Liu, M., Zhang, Z. & Xing, E. Roles of miRNA dysregulation in the pathogenesis of multiple myeloma, Cancer Gene Therapy 28, 1256–1268 doi.org/10.1038/s41417-020-00291-4 (2021)

9. van Rooij, E. and Kauppinen, S. Development of microRNA therapeutics is coming of age. EMBO Mol. Med. 6, 851–864. 10.1016/j.tig.2022.02.006a (2014).

10. Schirle, N. T., Sheu-Gruttadauria, J. & MacRae, I. J. Structural basis for microRNA targeting. Science 346, 608–613, doi:10.1126/science.1258040 (2014).

11. Chandradoss, S. D., Schirle, N. T., Szczepaniak, M., MacRae, I. J. & Joo, C. A Dynamic Search Process Underlies MicroRNA Targeting. Cell 162, 96–107, doi:10.1016/j.cell.2015.06.032 (2015).

12. Jo, M. H. et al. Human Argonaute 2 Has Diverse Reaction Pathways on Target RNAs. Mol Cell 59, 117–124, doi:10.1016/j.molcel.2015.04.027 (2015).

13. Baronti, L. et al. Base-pair conformational switch modulates miR-34a targeting of Sirt1 mRNA. Nature 583, 139-+, doi:10.1038/s41586-020-2336-3 (2020).

14. Han, J., Mendell, J. T. MicroRNA turnover: a tale of tailing, trimming, and targets, Trends in Biochemical Sciences, 1, 26-39, 10.1016/j.tibs.2022.06.005 (2023)

15. Helwak, A., Kudla, G., Dudnakova, T. & Tollervey, D. Mapping the Human miRNA Interactome by CLASH Reveals Frequent Noncanonical Binding. Cell 153, 654–665, doi:10.1016/j.cell.2013.03.043 (2013).

16. Chipman, L. B. & Pasquinelli, A. E. miRNA Targeting: Growing beyond the Seed. Trends Genet 35, 215–222, doi:10.1016/j.tig.2018.12.005 (2019).

17. Zhang, K. et al. A novel class of microRNA-recognition elements that function only within open reading frames. Nat Struct Mol Biol 25, 1019-+, doi:10.1038/s41594-018-0136-3 (2018).

18. Jame-Chenarboo, F; Ng, H. H; Macdonald, D; Mahal, L. K High-Throughput Analysis Reveals miRNA Upregulating α-2,6-Sialic Acid through Direct miRNA–mRNA Interactions. ACS Cent Sci, 8, 1527–1536 doi.org/10.1021/acscentsci.2c00748 (2022)

19. Jain, A. et al. Probing cellular protein complexes using single-molecule pull-down. Nature 473, 484–U322, doi:10.1038/nature10016 (2011)

20. Johnson-Buck, A. et al. Kinetic fingerprinting to identify and count single nucleic acids. Nat Biotechnol 33, 730–732, doi:10.1038/nbt.3246 (2015).

21. Bautista-Sanchez, D., Arriaga-Canon, C., Pedroza-Torres, A., De La Rosa-Velazquez, I.A., Gonzalez-Barrios, R., Contreras-Espinosa, L., Montiel-Manríquez, R., CastroHernandez, C., Fragoso-Ontiveros, V., Alvarez-Gomez, R.M., Herrera, L.A., 2020.The promising role of miR-21 as a cancer biomarker and its importance in RNAbased therapeutics. Molecular Therapy: Nucleic Acids, 20, 409–420, 10.1016/j.omtn.2020.03.003. (2020).

22. Hashemi M, Mirdamadi MSA, Talebi Y, Khaniabad N, Banaei G, Daneii P, et al. Pre-clinical and clinical importance of miR-21 in human cancers: tumorigenesis, therapy response, delivery approaches and targeting agents. Pharmacol Res, 187, 106568. 10.1016/j.phrs.2022.106568 (2023).

23. Geekiyanagea, H. Rayatpishehb, S. Wohlschlegel, J. A. Brown Jr, R and Ambros, V., Extracellular microRNAs in human circulation are associated with miRISC complexes that are accessible to anti-AGO2 antibody and can bind target mimic oligonucleotides, Proc. Natl. Acad. Sci 117, 39, 24213–24223 doi/10.1073/pnas.2008323117 (2020).

24. Ho, V. Baker, J. R. Willison, K. R. Barnes, P. J. Donnelly, L. E. and Klug, D. R. Single cell quantification of microRNA from small numbers of non-invasively sampled primary human cells, Communication Biology, 6, 458, doi.org/10.1038/s42003-023-04845-8 (2023).

25. Salomon, W. E., Jolly, S. M., Moore, M. J., Zamore, P. D. & Serebrov, V. Single-Molecule Imaging Reveals that Argonaute Reshapes the Binding Properties of Its Nucleic Acid Guides. Cell 162, 84–95, doi:10.1016/j.cell.2015.06.029 (2015).

26. Becker, W. R. et al. High-Throughput Analysis Reveals Rules for Target RNA Binding and Cleavage by AGO2. Mol Cell 75, 741-+, doi:10.1016/j.molcel.2019.06.012 (2019).

27. Brennecke, J. Stark, A. Russell, R. B. Cohen, S. M. Principles of MicroRNA–Target Recognition, PLoS Biology, 3, e85, doi.org/10.1371/journal.pbio.0030085 (2005).

28. Wee, L.M., Flores-Jasso, C.F., Salomon, W.E., and Zamore, P.D. (2012). Argonaute divides its RNA guide into domains with distinct functions and RNAbinding properties. Cell 151, 1055–1067.

29. Brennecke, J., Stark, A., Russell, R.B., and Cohen, S.M. (2005). Principles of microRNA-target recognition. PLoS Biol. 3, e85.

30. Lewis, B.P., Burge, C.B., and Bartel, D.P. (2005). Conserved seed pairing, often flanked by adenosines, indicates that thousands of human genes are microRNA targets. Cell 120, 15–20.

31. Grimson, A., Farh, K.K., Johnston, W.K., Garrett-Engele, P., Lim, L.P., and Bartel, D.P. (2007). MicroRNA targeting specificity in mammals: determinants beyond seed pairing. Mol. Cell 27, 91– 105.

32. Eden, U. T. & Kramer, M. A. Drawing inferences from Fano factor calculations. J Neurosci Meth 190, 149–152, doi:10.1016/j.jneumeth.2010.04.012 (2010).

33. Gourevitch, B. & Eggermont, J. J. A nonparametric approach for detection of bursts in spike trains. J Neurosci Meth 160, 349–358, doi:10.1016/j.jneumeth.2006.09.024 (2007).

34. Rinaldi, A. J., Lund, P. E., Blanco, M. R. & Walter, N. G. The Shine-Dalgarno sequence of riboswitch-regulated single mRNAs shows ligand-dependent accessibility bursts. Nat Commun 7, doi:ARTN 8976 10.1038/ncomms9976 (2016).

35. Klum, S.M., Chandradoss, S.D., Schirle, N.T., Joo, C., and MacRae, I.J. (2018). Helix-7 in Argonaute2 shapes the microRNA seed region for rapid target recognition. EMBO J. 37, 75–88. DOI 10.15252/embj.201796474.

36. Sheu-Gruttadauria, J., Xiao, Y., Gebert, L. F. R. & MacRae, I. J. Beyond the seed: structural basis for supplementary microRNA targeting by human Argonaute2. Embo J 38, doi:ARTN e101153 10.15252/embj.2018101153 (2019).

37. Sheu-Gruttadauria, J. et al. Structural Basis for Target-Directed MicroRNA Degradation. Mol Cell 75, 1243–1255 e1247, doi:10.1016/j.molcel.2019.06.019 (2019).

38. Willkomm, S. et al. Single-molecule FRET uncovers hidden conformations and dynamics of human Argonaute 2. Nat Commun 13, doi:ARTN 3825 10.1038/s41467-022-31480-4 (2022).

39. Welty, R., Schmidt, A. and Walter, N.G. (2023) Probing transient riboswitch structures *via* single molecule accessibility analysis. Methods Mol. Biol. 2568, 37–51, DOI: 10.1021/acschembio.3c00435.

40. Sheng, G. et al. Structure-based cleavage mechanism of Thermus thermophilus Argonaute DNA guide strand-mediated DNA target cleavage. Proc. Natl Acad Sci USA 111, 652–657, doi:10.1073/pnas.1321032111 (2014).

41. Wang, Y. et al. Nucleation, propagation and cleavage of target RNAs in Ago silencing complexes. Nature 461, 754–761, doi:10.1038/nature08434 (2009).

42. Wang, P. Y. Bartel, D. P. The guide-RNA sequence dictates the slicing kinetics and conformational dynamics of the Argonaute silencing complex, Molecular Cell, 84, 2918–2934 doi.org/10.1016/j.molcel.2024.06.026 (2024).

43. Khanna, K., Mandal, S., Blanchard, A. T., Tewari, M., Buck, A. J., Walter, N. G.,Rapid kinetic fingerprinting of single nucleic acid molecules by a FRET-based dynamic nanosensor. Biosensors and Bioelectronics 190, 113433, doi.org/10.1016/j.bios.2021.113433 (2021).

44. Walter, N G., Huang, C. Y., Manzo, A. J., & Sobhy, M. A., Do-it-yourself guide: how to use the modern single-molecule toolkit. Nat. Method. 5, 475, DOI:10.1038/NMETH.1215 (2008).

