## Supplementary Information for "A unifying model for microRNA-guided silencing of messenger RNAs"

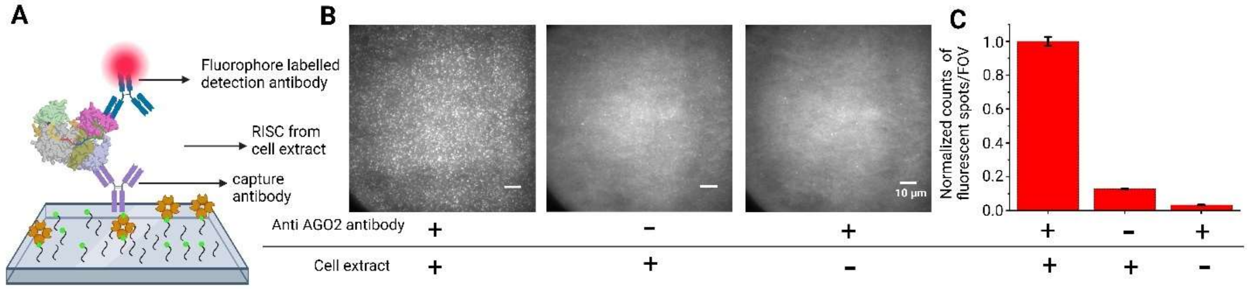


**Figure S1. Confirming capture of Ago2 from cell extract. Related to Figure 1**

(A) Schematic for detection of immunoprecipitated miRISC from cell extract using Cy5 labelled detection antibody. (B) Single molecule fluorescence microscope images of a field of view (FOV- 100 μM X 100 μM) showing thousands of bright puncta due to binding of Cy5 labelled detection antibody to miRISC (left image). Absence of either Ago2 antibody (middle image) or cell extract (right image) drastically reduces the number of fluorescent spots on the surface, indicating specific capture and detection of Ago2 by anti Ago2 antibodies from HeLa cell extract. The scale bar is indicated at the corner of each image. (C) Relative quantification of fluorescent spots/FOV on the surface in absence and presence of cell extract and capture anti Ago2 antibody. The number of fluorescent spots drops by 10-15 times in control experiments where either the cell extract or anti Ago2 antibody is absent. Values shown are the mean of independent duplicates ±SEM.

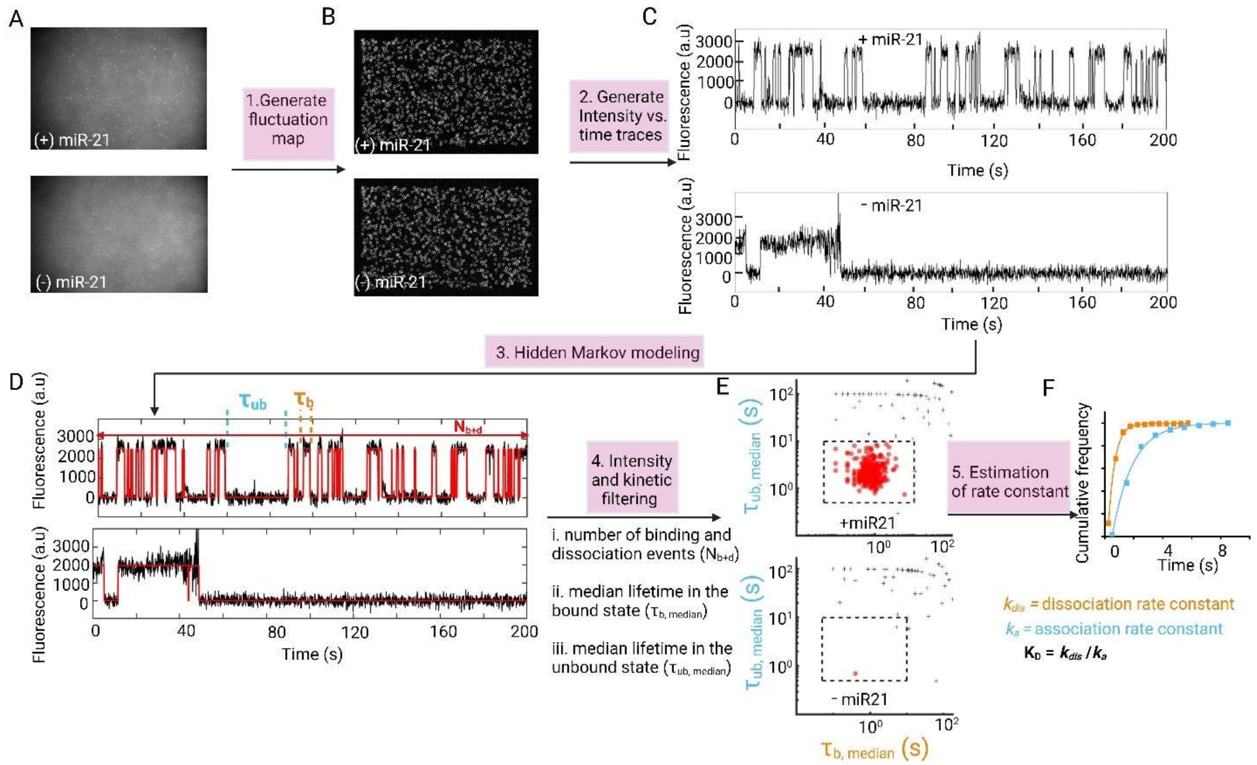


**Figure S2 SiMKEPS Data analysis pipeline. Related to Figure 1**

(A) Single-frame images of representative fields of view (FOV) from TIRF microscopy. (B) Intensity fluctuation maps of the fields of view shown in (A). Grey circles indicate positions of local maxima in the fluctuation map, from which candidate ROIs are identified for further analysis to generate intensity vs. time traces. (C) Representative intensity vs. time traces generated from the ROIs identified in (B). (D) Hidden Markov modelling (HMM) idealization (red lines) for each intensity vs. time trace. Bound and unbound-state dwell times ($\tau_{b,median}$ and $\tau_{ub,median}$ respectively) are indicated by the orange and blue horizontal line segments above the idealization. Number of binding and dissociation events ($N_{b+d}$) are shown as the vertical red lines and indicates how many times the fluorescent probe undergoes association and dissociation with a single miR-21 RISC (E) Candidates in the positive (orange circles) and negative (blue squares) controls for miR-21 RISC are well separated by thresholds of $N_{b+d} > 20$ and $\tau_{bound} > 0.5$ s (black dashed lines), permitting discrimination of specific and nonspecific binding at the single-molecule level. Data are pre-filtered for signal-to-noise > 2.5 and intensity > 200. (F) Fitting an exponential cumulative distribution function (CDF) to the dwell times in the bound state ($t_{b,\ median}$) and unbound state ($t_{ub,\ median}$) of individual kinetically filtered time traces provides association rate constant ($k_a$), dissociation rate constant ($k_{dis}$) and affinity ($K_D = k_{dis} / k_a$) of an interaction.

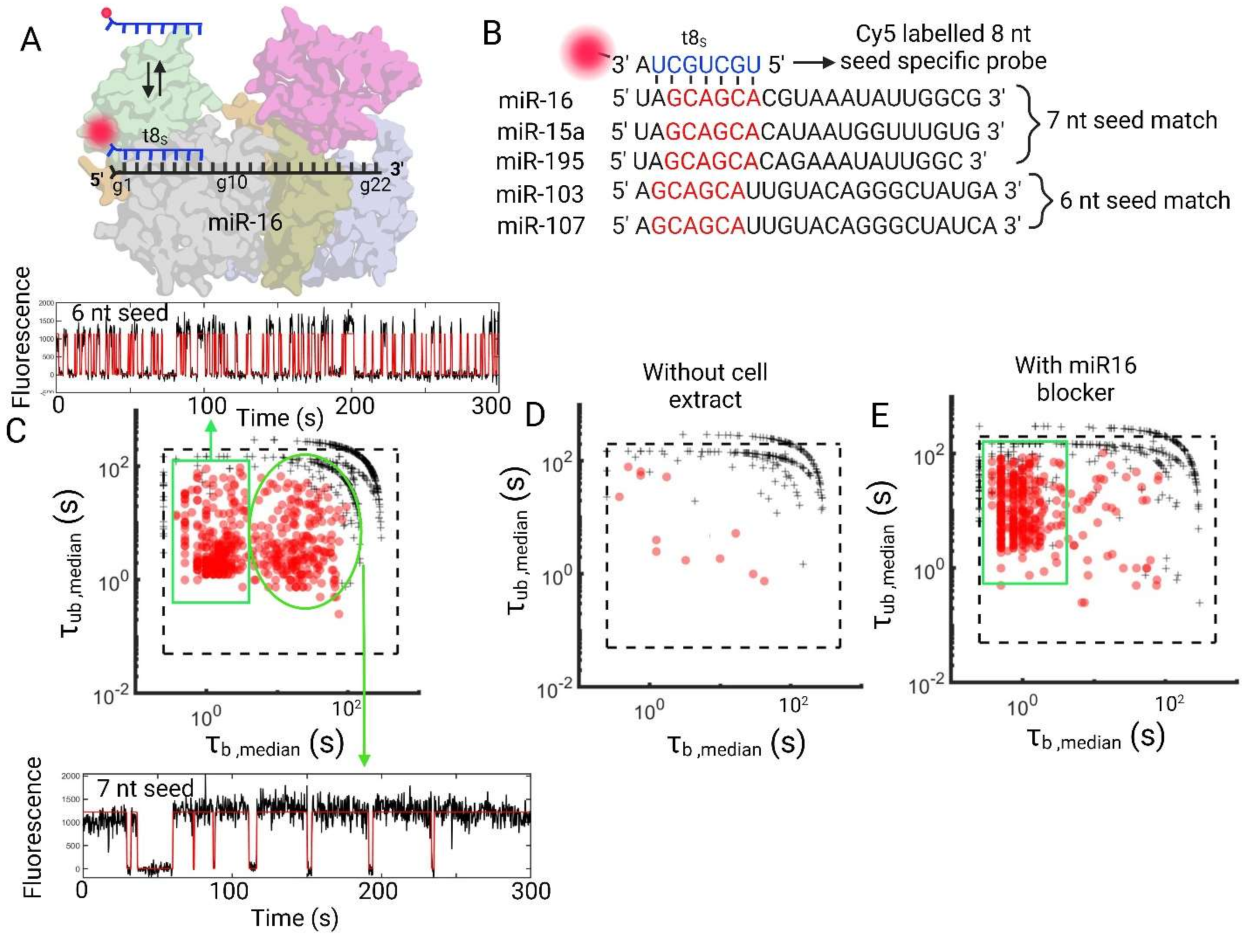


**Figure S3. Detection of miR-16 and its family members. Related to Figure 1**

(A) Schematic showing interaction of target mimic probe ($t8_{7S}$) of miR-16, with miR-16 RISC complexes. (B) Sequences of miR-16 seed family members (miR15a and miR195) and miR-103/miR-107 family members. miR-103/miR-107 has 6 nucleotide seed matches (shown in red) to miR-16 seed family members. The sequence of 8 nt target mimic ($t8_{7S}$) has shown in blue. (C) SiMKEPS experiment show interaction of $t8_{7S}$ with miRISC, immunoprecipitated from cell extract, generates two distinct clusters of interaction in the scatter plot of $\tau_{bound}$ vs $\tau_{unbound}$ dwell time. One of those interaction is ~20 times more stable than the other interaction. The kinetic time traces corresponding to each cluster are shown. The relatively weaker interaction is characterized as interaction of $t8_{7S}$ with 6 nt seed match guide miRNA (miR-103/miR-107 family members) and relatively stable interaction has been characterized as interaction with 7 nt seed match guide miRNAs (miR-16 family members). (D) In absence of cell extract or in presence of only Ago2, devoid of any miRNAs, no specific interaction was observed. (E) Addition of miR-16 specific blocker removes one of this cluster having relatively stable interaction with $t8_{7S}$. However, the relatively weak interaction remains, indicating miR-16 blocker cannot efficiently blocks the miR-103/miR-107 family members.

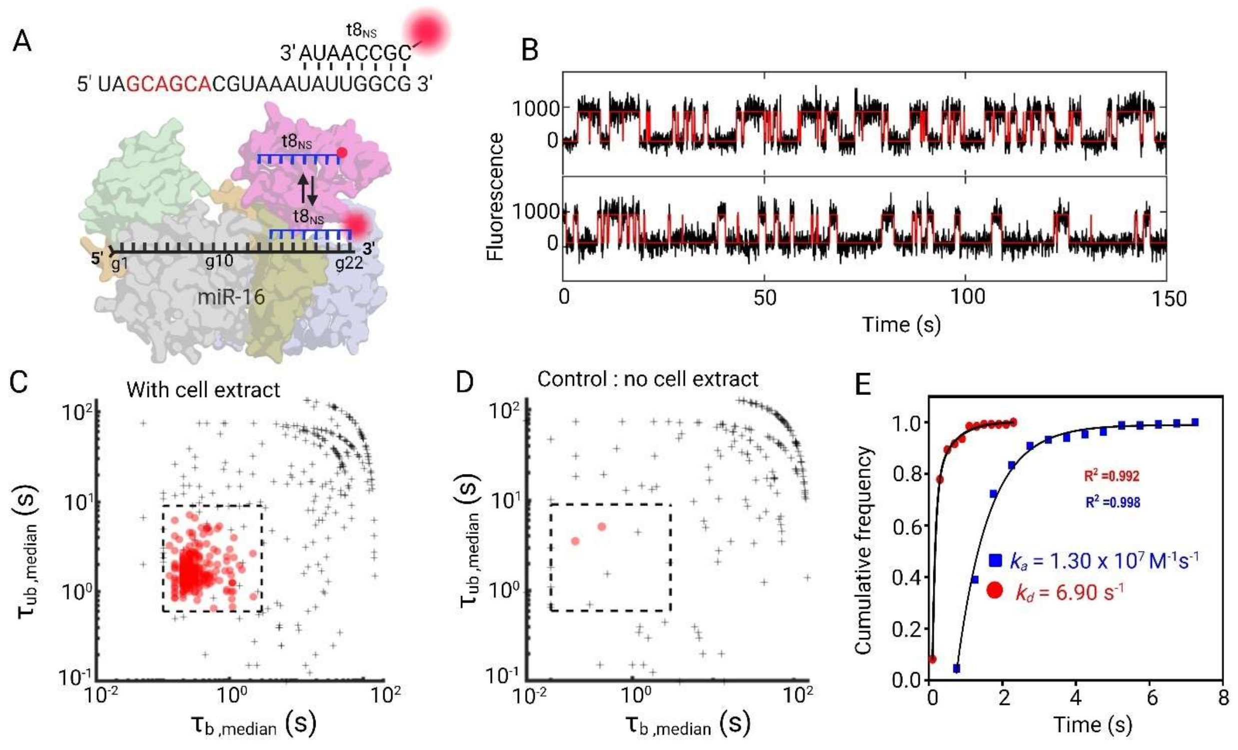


**Figure S4. Detection of miR-16. Related to Figure 1**

(A) Schematic showing interaction of target mimic probe ($t8_{NS}$) of miR-16, with miR-16 RISC. (B) Example fluorescence intensity time trajectories showing interaction of $t8_{NS}$ probe with miR-16 RISC. (C) SiMKEPS experiment show interaction of $t8_{7S}$ with miRISC, immunoprecipitated from cell extract, generates clusters of interaction in the scatter plot of $\tau_{bound}$ vs $\tau_{unbound}$ dwell time. (D) In absence of cell extract or in presence of only Ago2, devoid of any miRNAs, no specific interaction was observed. (E) Association and dissociation rate constant was estimated from exponential fitting of cumulative distribution of $\tau_{unbound}$ and $\tau_{bound}$ and dwell times respectively.

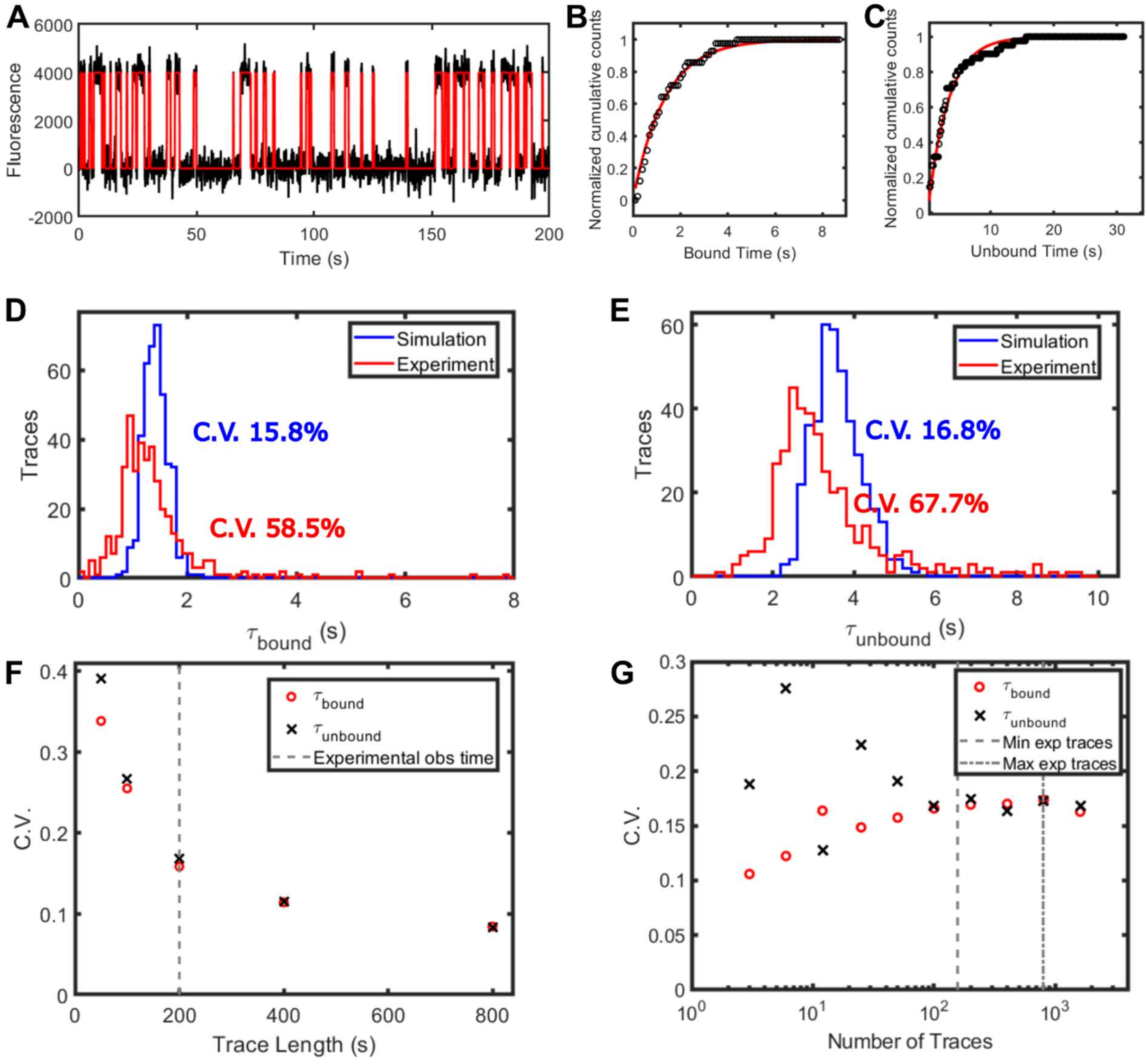


**Figure S5. Data simulation show heterogeneity in the target-miRISC interaction. Related to Figures 1 and 3**

(A) Representative simulated single-molecule intensity trace with kinetics similar to the binding of $t8_{7s}$ to miR-21 RISC. The black line represents the raw intensity signal with Gaussian noise added to simulate experimental noise, while the red line represents a hidden Markov modeling idealization. (B-C) Single-exponential fits to cumulative distribution functions of bound- (B) and unbound-state (C) dwell times for the simulated intensity trace of a single miRISC complex in (A). (D, E) Estimated bound- (D) and unbound-state (E) lifetimes derived from single-exponential fits of cumulative distributions of dwell times from either simulated (blue, *N*=400) or experimental (red, *N*=392) traces of $t8_{7s}$ binding. Coefficients of variation (C.V.) for each distribution are shown. (F-G) Dependence of the C.V. of bound- and unbound-state lifetime estimates from single-exponential fitting of single simulated traces as a function of trace length with a constant number of *N* = 400 traces (F) or as a function of the number of traces with a constant trace length *l* = 2000 frames (200 s) (G). Since traces are simulated to reflect a single rate constant each for binding and dissociation, (F) and (G) give an estimate of the maximum achievable precision for single-molecule kinetic measurements with SiMKEPS in the presence of completely homogeneous behavior. The vertical dashed line in (F) corresponds to the experimental measurement time of 200 s, and the vertical dashed lines in (G) correspond to the minimum and maximum number of traces per experimental condition (156 and 792, respectively). Under these experimental conditions, statistical error is not expected to result in C.V.s greater than 0.2 (F), and estimates of C.V. are likely to be stable and accurate (G).

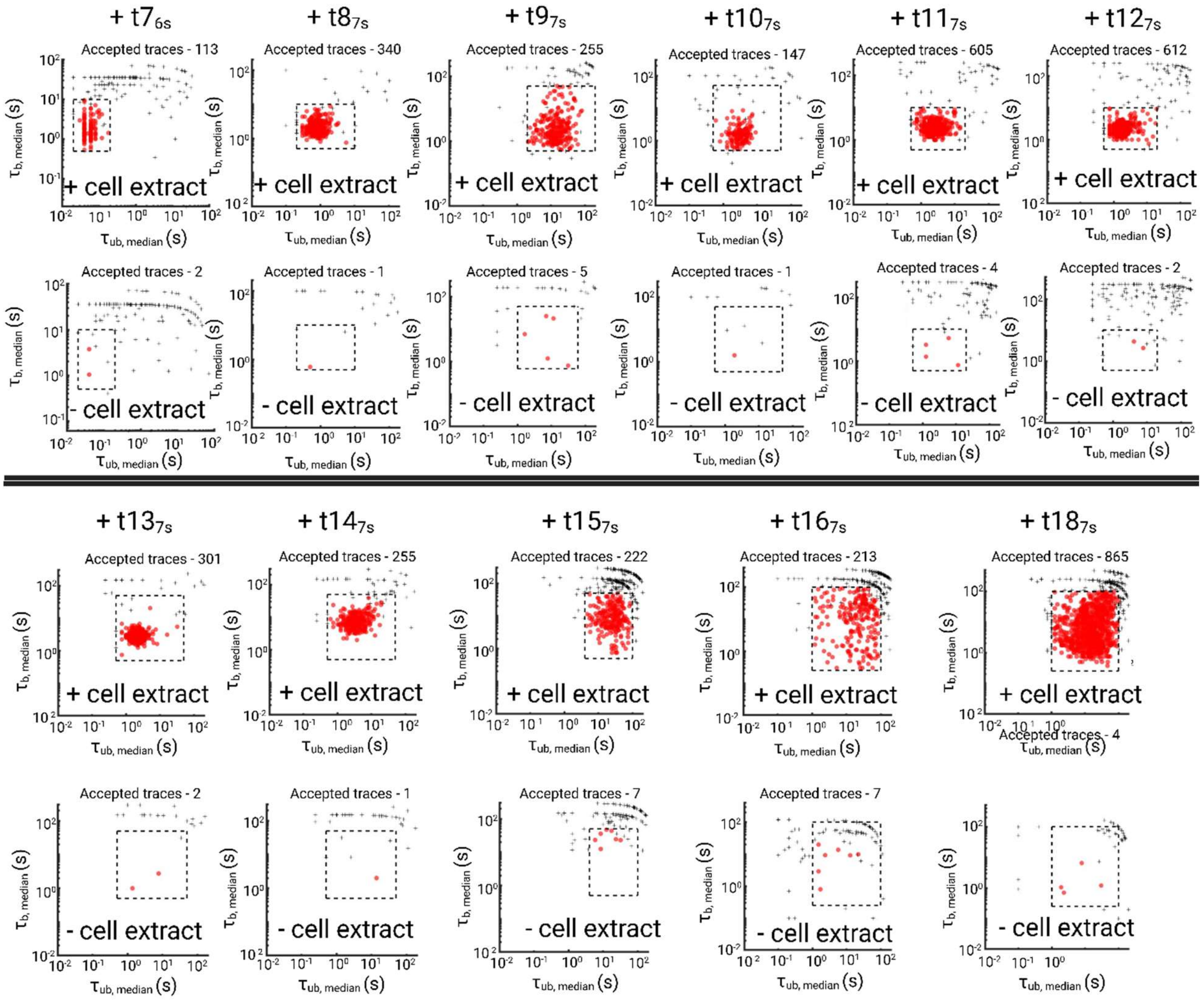


**Figure S6. Kinetic filtering can separate specific from non-specific interaction. Related to Figure 2**

Interaction of surface captured miR-21 RISC with varying length ($t7_{6S}$ – $t18_{7S}$) of 3’ Cy5 labelled target mimic. Scatter plot distribution of $\tau_{b,median}$ and $\tau_{ub,median}$ show kinetic filtering (shown as black dotted line In the Figure and tabulated in Table S1) can separates the specific interaction from non-specific interaction (comparison of ‘+ cell extract’ and ‘– cell extract’).

**Table S1. related to Figure 2**

Kinetic filtering parameters to separate specific interaction of miR-21 RISC with target mimics has been tabulated below.

| | Intensity threshold | Signal/Noise (S/N) | $N_{b+d}$ | $\tau_b$ (min) | $\tau_b$(max) | $\tau_{ub}$ (min) | $\tau_{ub}$ (max) |
|---|---|---|---|---|---|---|---|
| $t7_{6s}$ | 200 | 3 | 10 | 0.02 | 0.2 | 0.5 | 10 |
| $t8_{7s}$ | 200 | 3 | 15 | 0.2 | 10 | 0.5 | 10 |
| $t9_{7s}$ | 200 | 3 | 4 | 2 | 200 | 0.5 | 50 |
| $t10_{7s}$ | 500 | 3 | 12 | 0.5 | 50 | 0.5 | 50 |
| $t11_{7s}$ | 500 | 3 | 15 | 0.5 | 20 | 0.5 | 10 |
| $t12_{7s}$ | 500 | 3 | 15 | 0.5 | 20 | 0.5 | 10 |
| $t13_{7s}$ | 500 | 3 | 8 | 0.5 | 50 | 0.5 | 50 |
| $t14_{7s}$ | 500 | 3 | 8 | 0.5 | 50 | 0.5 | 50 |
| $t15_{7s}$ | 200 | 3 | 4 | 4 | 100 | 0.5 | 50 |
| $t16_{7s}$ | 200 | 3 | 4 | 1 | 100 | 0.25 | 100 |
| $t18_{7s}$ | 200 | 3 | 4 | 1 | 100 | 0.25 | 100 |

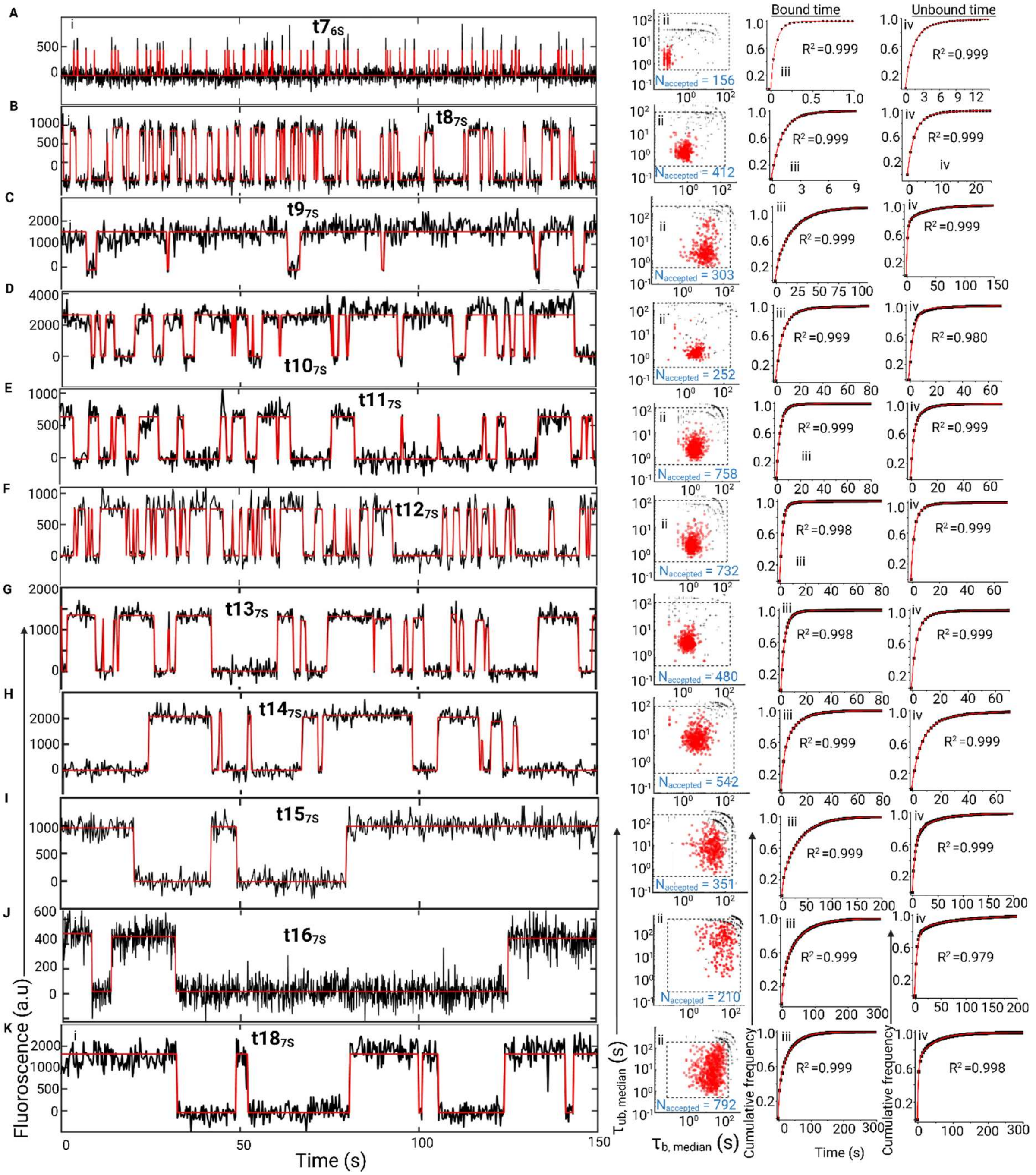


**Figure S7. Interaction kinetics of different length probe with miRISC. Related to Figure 2**

Interaction of surface captured miR-21 RISC with varying length of 3' Cy5 labelled target mimic (A; t7$_{6S}$, B; t8$_{7S}$, C; t9$_{7S}$, D; t10$_{7S}$, E; t11$_{7S}$, F; t12$_{7S}$, G; t13$_{7S}$, H; t14$_{7S}$ I; t15$_{7S}$, J;

t16$_{7S}$, K; t18$_{7S}$). i. Intensity vs time trajectories of interaction between miR-21 RISC and seed probe of different lengths ii. Scatter plots of bound and unbound time distribution of different target mimic probe. iii. Fitting an exponential cumulative distribution function (CDF) to the dwell times in the bound state ($\tau_b$) provides dissociation rate constant ($k_{dis}$) and iv. Fitting CDF to the dwell times in the unbound state ($\tau_{ub}$) provides association rate constant ($k_a$).

**Table S2. Kinetic parameters extracted from SiMKEPS analysis. related to Figure 2**

Sequences of seed (S) probe used in this study. Corresponding association rate constant ($k_a$) dissociation rate constant ($k_d$) and affinity ($K_D$) of interaction between seed probe and miR-21 guide strand has been tabulated.

| Probe [illegible]esignation | Sequence of probes | [illegible]ssociation rate ([illegible]$_a$ x $10^7$) $M^{-1}s^{-1}$ | [illegible]issociation rate ($k_{dis}$) $s^{-1}$ | [illegible] = $k_{dis}/k_a$ nM |
|---|---|---|---|---|
| $t7_{6S}$ | 5’ UAAGCUA 3’Cy5 | 0.89 ± 0.053 | 5.658 ± 0.941 | [illegible]35 ± 69 |
| $t8_{7S}$ | 5’ AUAAGCUA 3’ Cy5 | 0.74 ± 0.061 | 0.625 ± 0.032 | [illegible]4 ± 12 |
| $t9_{7S}$ | 5’ GAUAAGCUA 3’ Cy5 | 0.12 ± 0.010 | 0.041 ± 0.008 | [illegible]4 ± 7 |
| $t10_{7S}$ | 5’ UGAUAAGCUA 3’ Cy5 | 0.42 ± 0.045 | 0.187 ± 0.035 | [illegible]5 ± 5 |
| $t11_{7S}$ | 5’ CUGAUAAGCUA 3’ Cy5 | 0.41 ± 0.043 | 0.281 ± 0.023 | [illegible]8 ± 5 |
| $t12_{7S}$ | 5’ UCUGAUAAGCUA 3’ Cy5 | 0.45 ± 0.035 | 0.390 ± 0.031 | [illegible]6 ± 6 |
| $t13_{7S}$ | 5’ GUCUGAUAAGCUA 3’ Cy5 | 0.31 ± 0.025 | 0.265 ± 0.026 | [illegible]5 ± 8 |
| $t14_{7S}$ | 5’ AGUCUGAUAAGCUA 3’ Cy5 | 0.16 ± 0.019 | 0.126 ± 0.007 | [illegible]9 ± 7 |
| $t15_{7S}$ | 5’ CAGUCUGAUAAGCUA 3’ Cy5 | 0.11 ± 0.020 | 0.034 ± 0.005 | [illegible]1 ± 8 |
| $t16_{7S}$ | 5’ UCAGUCUGAUAAGCUA 3’ Cy5 | [illegible].078 ± 0.018 | 0.021 ± 0.008 | [illegible]7 ± 10 |
| $t18_{7S}$ | [illegible]AUCAGUCUGAUAAGCUA 3’ Cy5 | [illegible].076 ± 0.016 | [illegible]015 ± 0.005 | [illegible]0 ± 9 |
| $t9_{7S}$ + 3A | 5’ AAAGAUAAGCUA 3’ Cy5 | 0.35 ± 0.035 | 0.263 ± 0.024 | [illegible]5 ± 8 |
| $t8_{7S}$ + 4A | 5’ AAAAAUAAGCUA 3’ Cy5 | 0.54 ± 0.083 | 0.471 ± 0.048 | [illegible] ± 8 |

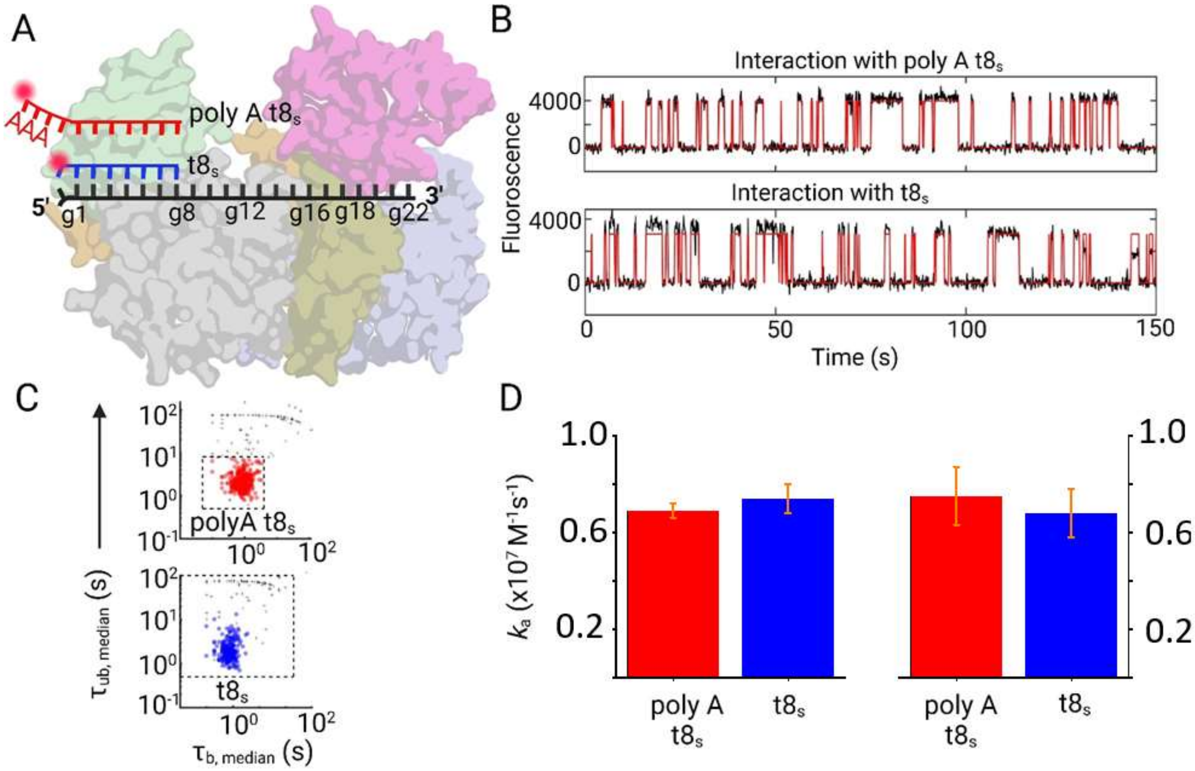


**Figure S8. Positional effect of fluorophore on seed interaction. Related to Figure 2**

(A) Schematic showing interaction of target mimic probe $t8_{7S}$ and poly A $t8_{7S}$ with miR-21 RISC complexes. Compared to $t8_{7S}$, 3' Cy5 probe is further distant from g1 nucleotide of guide strand by three adenosine nucleotide of poly A $t8_{7S}$. (B) Intensity vs time trajectories of interaction of poly A $t8_{7S}$ (upper trace) and $t8_{7S}$ (lower trace) with miR-21 RISC. (C) Scatter plots of bound and unbound time distribution of poly A $t8_{7S}$ (upper plot) and $t8_{7S}$ (lower plot). (D) Association and dissociation rate calculation of poly A $t8_{7S}$ and $t8_{7S}$ indicate negligible effect of fluorophore on interaction kinetics of seed probe with miR-21 RISC. Mean of two independent replicates: ±SEM.

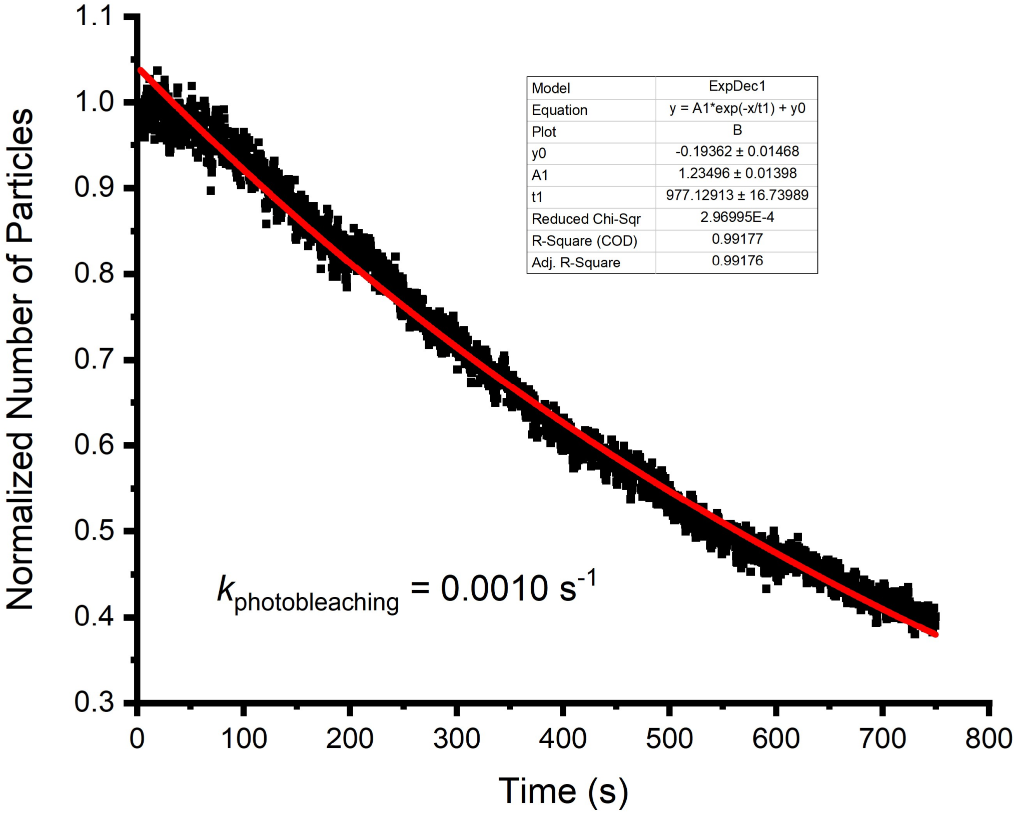


**Figure S9. Quantification of Cy5 photobleaching. Related to Figure 2**

Rate constant of Cy5 photobleaching under the illumination conditions of our experiments (~4 mW laser output power), provide lower-bound estimates of the photobleaching time constant ($k_{photobleaching}$) for each fluorophore.

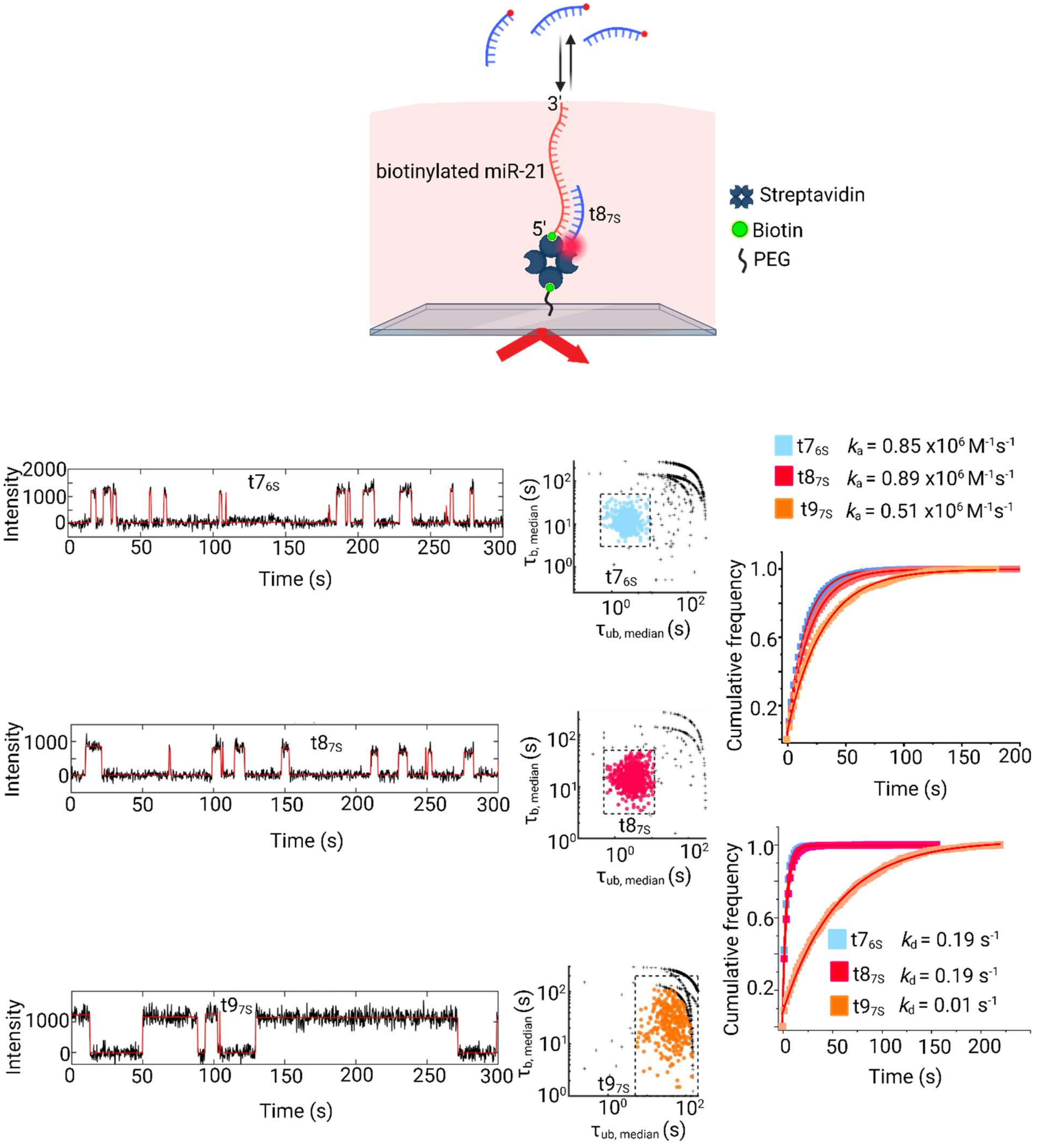


**Figure S10. Kinetics of protein-free miRNA-21 with target mimic. Related to Figure 2**

Interaction of $t7_{6S}$, $t8_{7S}$ and $t9_{7S}$ probe with protein-free biotinylated miR-21. (A) Intensity vs time trajectories of $t7_{S}$ (upper trace), $t8_{7S}$ (middle trace) and $t9_{7S}$ (lower trace) with protein-free biotinylated miR-21. (B) Scatter plots of the median $\tau_{bound}$ and $\tau_{unbound}$ dwell times for all intensity-versus-time trajectories within a single field of view of $t7_{S}$ (upper panel) and $t8_{7S}$ (middle panel) and $t9_{7S}$ (lower trace). (C) Cumulative distribution fitting of unbound time of $t7_{S}$, $t8_{7S}$ and $t9_{7S}$; corresponding association rate are shown with respective color (D) Cumulative distribution fitting of bound time of $t7_{S}$, $t8_{7S}$ and $t9_{7S}$. corresponding dissociation rate are shown with respective color.

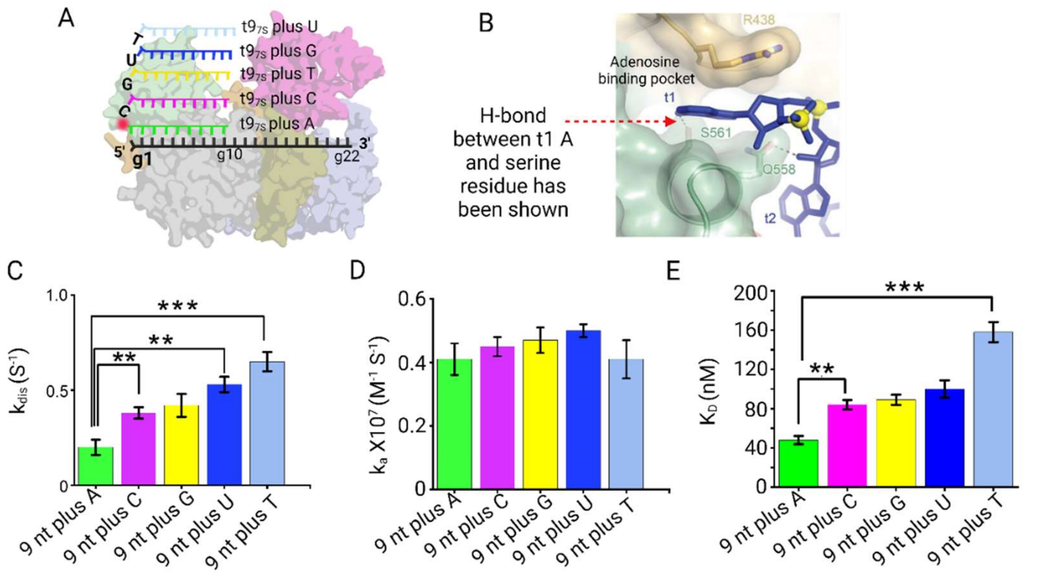


**Figure S11. Effect of Adenosine nucleotide at t1 position of target. Related to Figure 2**

(A) Schematic showing the sequences of different 10nt Seed probe; Adenine (9 nt plus A - green), Cytosine (9 nt plus C - pink), Guanine (9 nt plus G - yellow), Uracil (9 nt plus U - blue) or Thymine (9 nt plus T – faded blue) at t1 position. and complementary to the guide miR-21RISC (B) Binding pocket for t1 Adenine between L2 and MID domains. H-bond between t1 Adenosine and serine 561 residue has been indicated by red broken arrow. (C) Dissociation rate constant ($K_{dis}$) of 10 nt probe ($t10_{7S}$) having either Adenine (9 nt plus A), Cytosine (9 nt plus C), Guanine (9 nt plus G), Uracil (9 nt plus U) or Thymine (9 nt plus T) at t1 position. Mean of three independent replicates: ±SEM (***$p < 0.01$, **$p < 0.05$, *$p < 0.1$) (D) Association rate constant ($K_a$) of 10 nt probe ($t10_{7S}$) having either Adenine, Cytosine, Guanine, Uracil or Thymine at t1 position. Mean of three independent replicates: ±SEM (E) Binding affinity (dissociation equilibrium constant, $K_D$) of target mimics having varying t1 nucleotide to miR-21 RISC. Mean of independent triplicates ± SEM. (***$p < 0.01$, **$p < 0.05$, *$p < 0.1$)

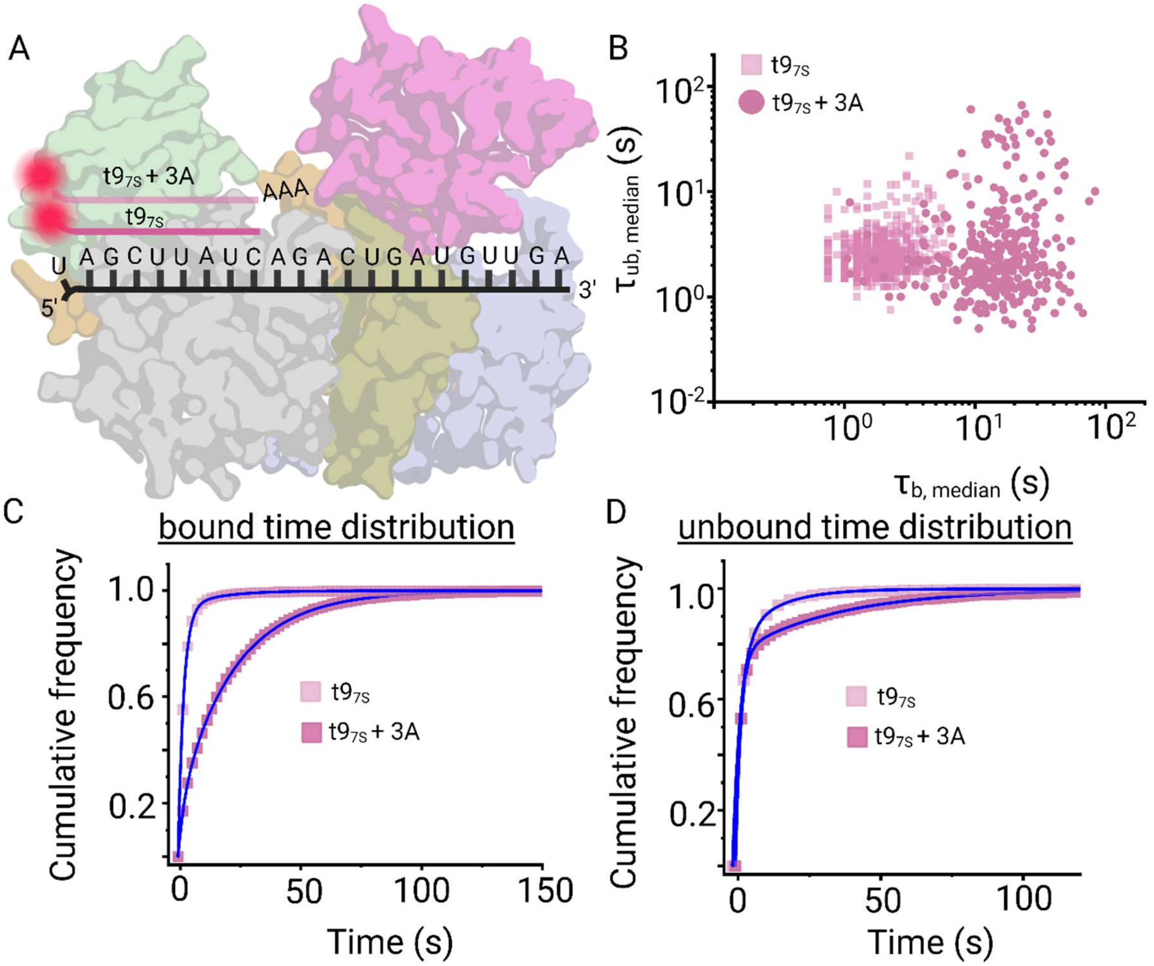


**Figure S12. Effect of Adenosine nucleotides at 5' position of t9$_{7S}$. Related to Figure 2**

(A) Schematic showing interaction of target mimic probe t9$_{7S}$ and t9$_{7S}$ + 3A having three adenosines at 5' of t9$_{7S}$ with guide miR-21. (B) Scatter plots of bound and unbound time for t9$_{7S}$ and t9$_{7S}$ + 3A. Cumulative distribution plot of unbound (C) and bound time (D) of t9$_{7S}$ and t9$_{7S}$ + 3A.

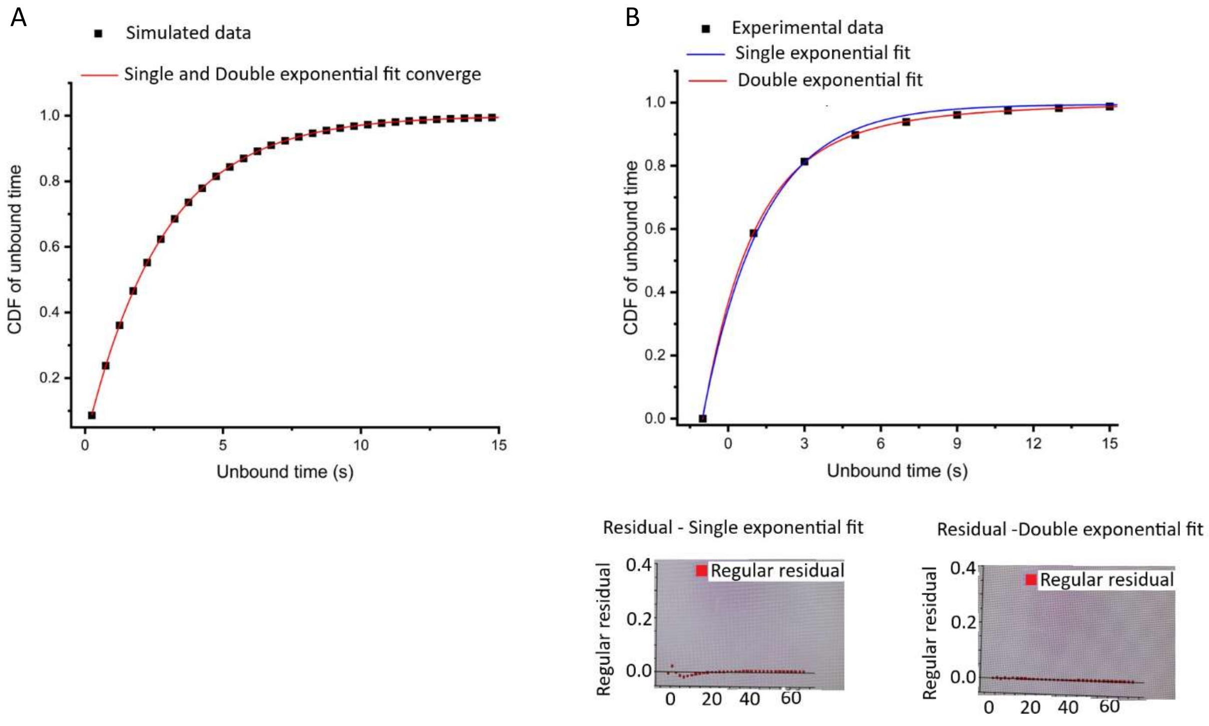


**Figure S13. Double-exponential fits better than single-exponential fitting of experimental data. Related to Figure 3**

Single and double-exponential fit of Simulated (A) and experimental data (B) for interaction between miRISC and $t8_{7S}$. (A) Single-exponential fit ($y = m\ e^{-x/\tau} + c$) and double-exponential fit ($y = m\ e^{-x/\tau 1} + n\ e^{-x/\tau 2} + c$) converge for Simulated data points (B) Single-exponential fit and double-exponential fit does not converge for experimental data points. Residual of fitting show that a double-exponential equation fits the data points better than single-exponential equation.

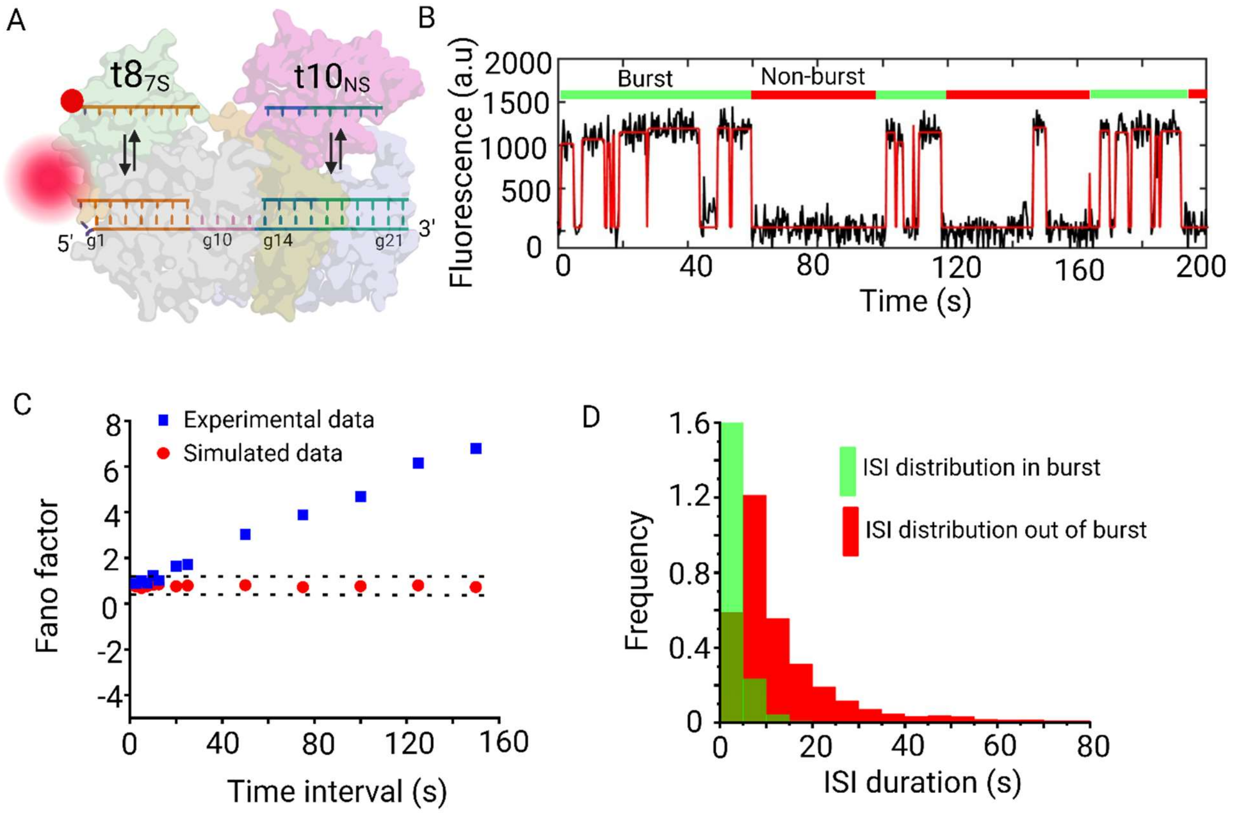


**Figure S14. Secondary structure formation of miR-21 is not a reason to explain bursting behavior in seed probe binding. Related to Figure 3**

(A) Effect of excess (1 μM) non-labeled 10 nt non-seed probe ($t10_{NS}$) on bursting behavior of 8 nt seed ($t8_{7S}$) probe. (B) Traces showing bursting in $t8_{7S}$ probe binding to miR-21 RISC even in presence of $t10_{NS}$. (C) Fano factor calculation and (D) inter spike interval (ISI) in burst and non-burst periods indicate that conformational change in miRISC still happen when the 3' NS region is blocked by the 10 nt NS probe ($t10_{NS}$)

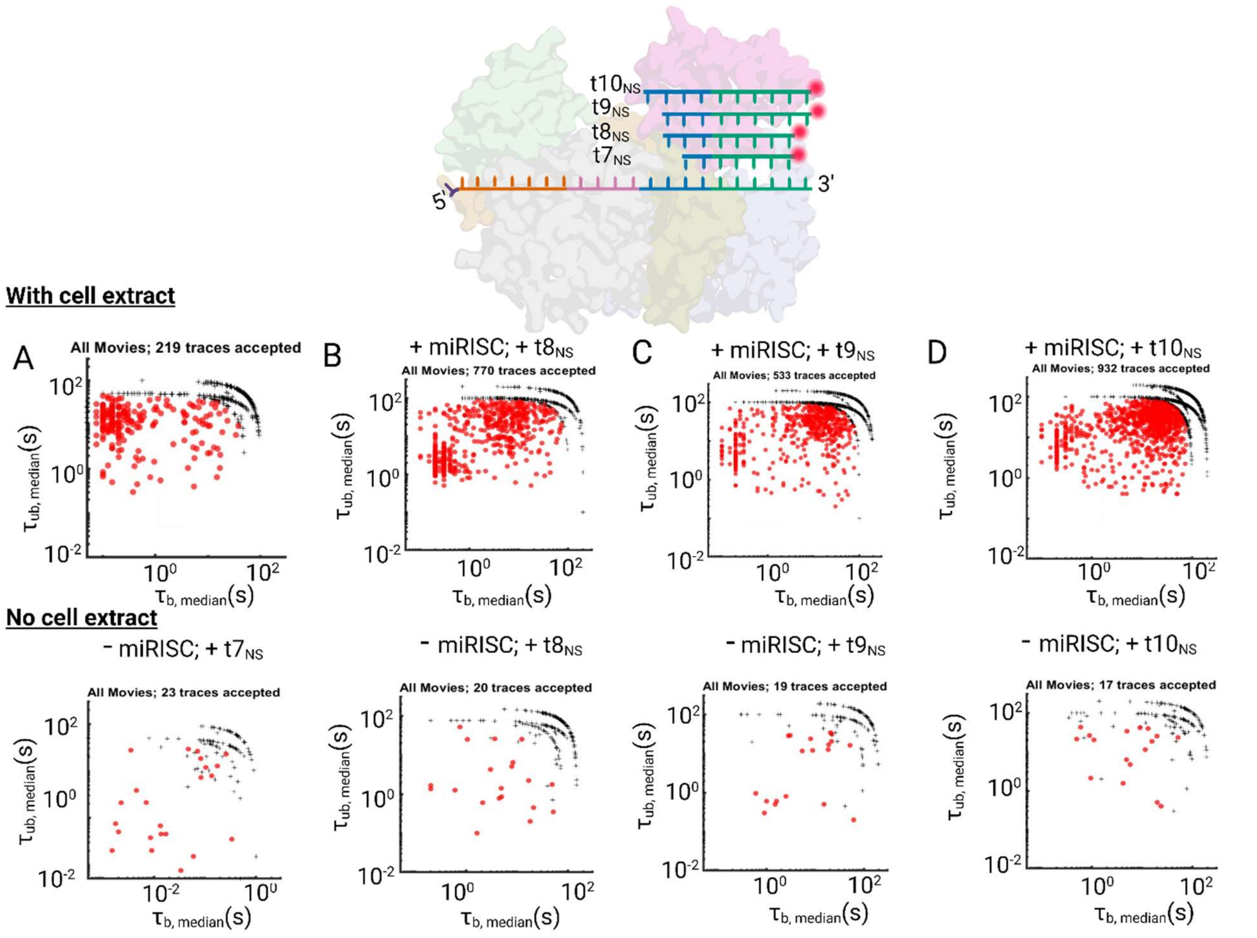


**Figure S15. Scatter plots of the median $\tau_{bound}$ and $\tau_{unbound}$ dwell times for all supplementary and 3' NS binding probe. Related to Figure 4**

Scatter plots of the median $\tau_{bound}$ and $\tau_{unbound}$ dwell times for all intensity-versus-time trajectories within a single field of view in the presence of Cy5-labeled target mimics of different length interacting with miR-21 RISC (with cell extract; upper scatter plots and no cell extract control; lower scatter plots). (A) $t7_{NS}$, (B) $t8_{NS}$, (C) $t9_{NS}$ and (D) $t10_{NS}$

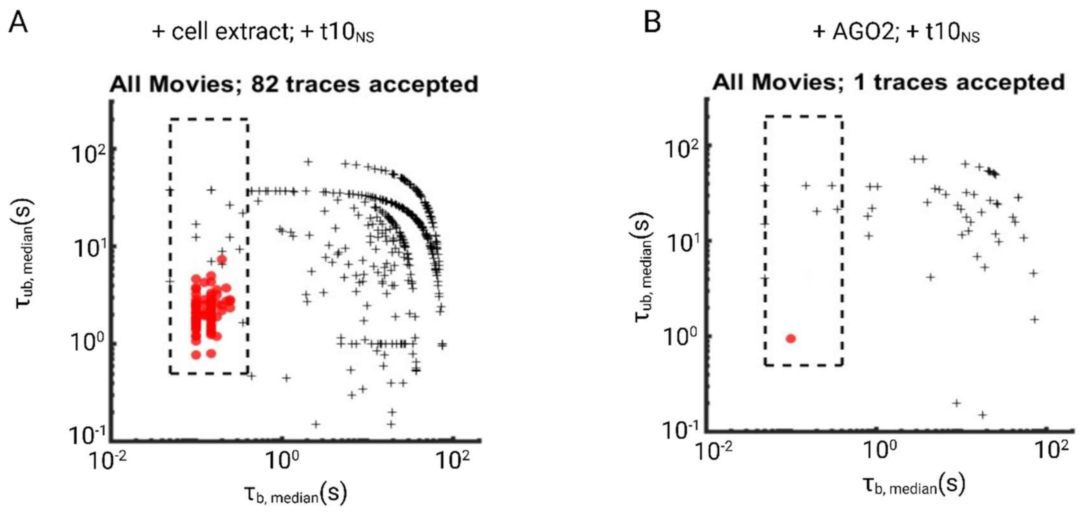


**Figure S16. Non-seed probe does not detectably interact with Ago2. Related to Figure 4**

Scatter plots of the median $\tau_{bound}$ and $\tau_{unbound}$ dwell times for all intensity-versus-time trajectories within a single field of view in the presence of Cy5-labeled $t10_{NS}$ target mimic interacting with miR-21 RISC (A) (with cell extract; left scatter plot) and (B) with purified AGO2 devoid of any miRNA (right scatter plot).

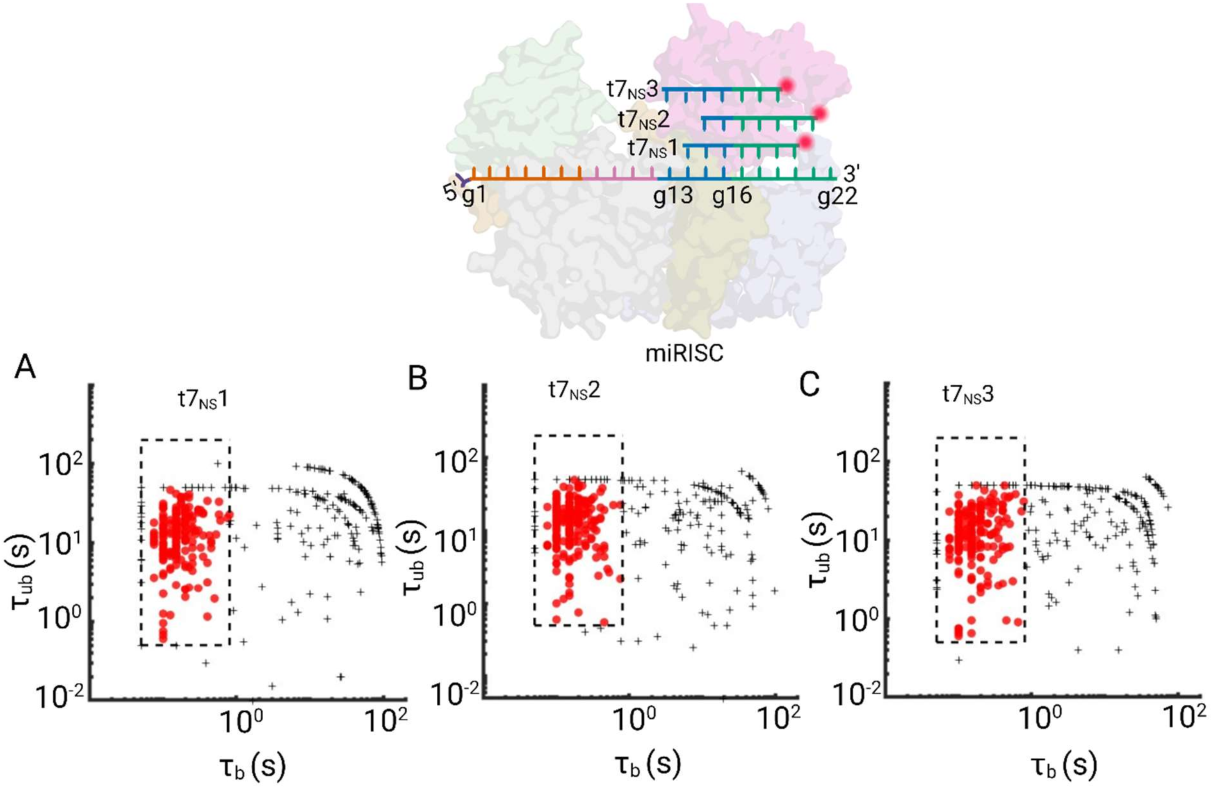


**Figure S17. Transient interaction kinetics of 7 nt NS probe ($t7_{NS}$) with miR-21 RISC is independent of its sequence. Related to Figure 4.**

(A, B, C) Scatter plots of the median $τ_{bound}$ and $τ_{unbound}$ dwell times for all intensity-versus-time trajectories within a single field of view for $t7_S$ having perfect base complimentary to different regions of miR-21 guide strand.

**Table S3. Related to Figure 4**.

Sequences of non-seed (NS) probe used in this study. Association rate constant ($k_a$) dissociation rate constant ($k_d$) and affinity ($K_D$) for transient interaction component.

| Probe designation | Sequences of probe | Association rate ($k_a$ x $10^7$) $M^{-1}s^{-1}$ | Dissociation rate ($k_{dis}$) $s^{-1}$ | $K_D = k_{dis}/k_a$ nM |
|---|---|---|---|---|
| $t7_{NS}$ (1) | Cy5 5' AACAUCA 3' | 0.18 ± 0.09 | 8.2 ± 0.35 | 4555.5 ± 232 |
| $t7_{NS}$ (2) | Cy5 5' CAACAUC 3' | 0.23 ± 0.05 | 9.8 ± 0.76 | 4260.86 ± 357 |
| $t7_{NS}$ (3) | Cy5 5' ACAUCAG 3' | 0.21 ± 0.05 | 9.6 ± 0.29 | 4571.42 ± 378 |
| $t8_{NS}$ | Cy5 5' CAACAUCA 3' | 1.05 ± 0.13 | 4.93 ± 0.17 | 469 ± 60 |
| $t9_{NS}$ | Cy5 5' UCAACAUCA 3' | 0.61 ± 0.06 | 3.92 ± 0.23 | 642 ± 90 |
| $t10_{NS}$ | Cy5 5' UCAACAUCAG 3' | 0.60 ± 0.05 | 4.50 ± 0.79 | 750 ± 92 |

**Table S4. Related to Figure 4**

Association rate constant ($k_a$) dissociation rate constant ($k_d$) and affinity ($K_D$) for long-lived binding component.

| Probe designation | Sequences of probe | Association rate ($k_a$ x $10^6$) $M^{-1}s^{-1}$ | Dissociation rate ($k_{dis}$) $s^{-1}$ | $K_D = k_{dis}/k_a$ nM |
|---|---|---|---|---|
| $t7_{NS}$ (1) | Cy5 5' AACAUCA 3' | 1.08 ± 0.09 | 0.48 ± 0.09 | 444 ± 112 |
| $t7_{NS}$ (2) | Cy5 5' CAACAUC 3' | 1.20 ± 0.21 | 0.53 ± 0.12 | 441 ± 89 |
| $t7_{NS}$ (3) | Cy5 5' ACAUCAG 3' | 0.71 ± 0.08 | 0.33 ± 0.08 | 464 ± 78 |
| $t8_{NS}$ | Cy5 5' CAACAUCA 3' | 0.75 ± 0.05 | 0.071 ± 0.023 | 94 ± 18 |
| $t9_{NS}$ | Cy5 5' UCAACAUCA 3' | 0.47 ± 0.09 | 0.046 ± 0.005 | 97 ± 27 |
| $t10_{NS}$ | Cy5 5' UCAACAUCAG 3' | 0.51 ± 0.08 | 0.01 ± 0.004 | 65 ± 19 |

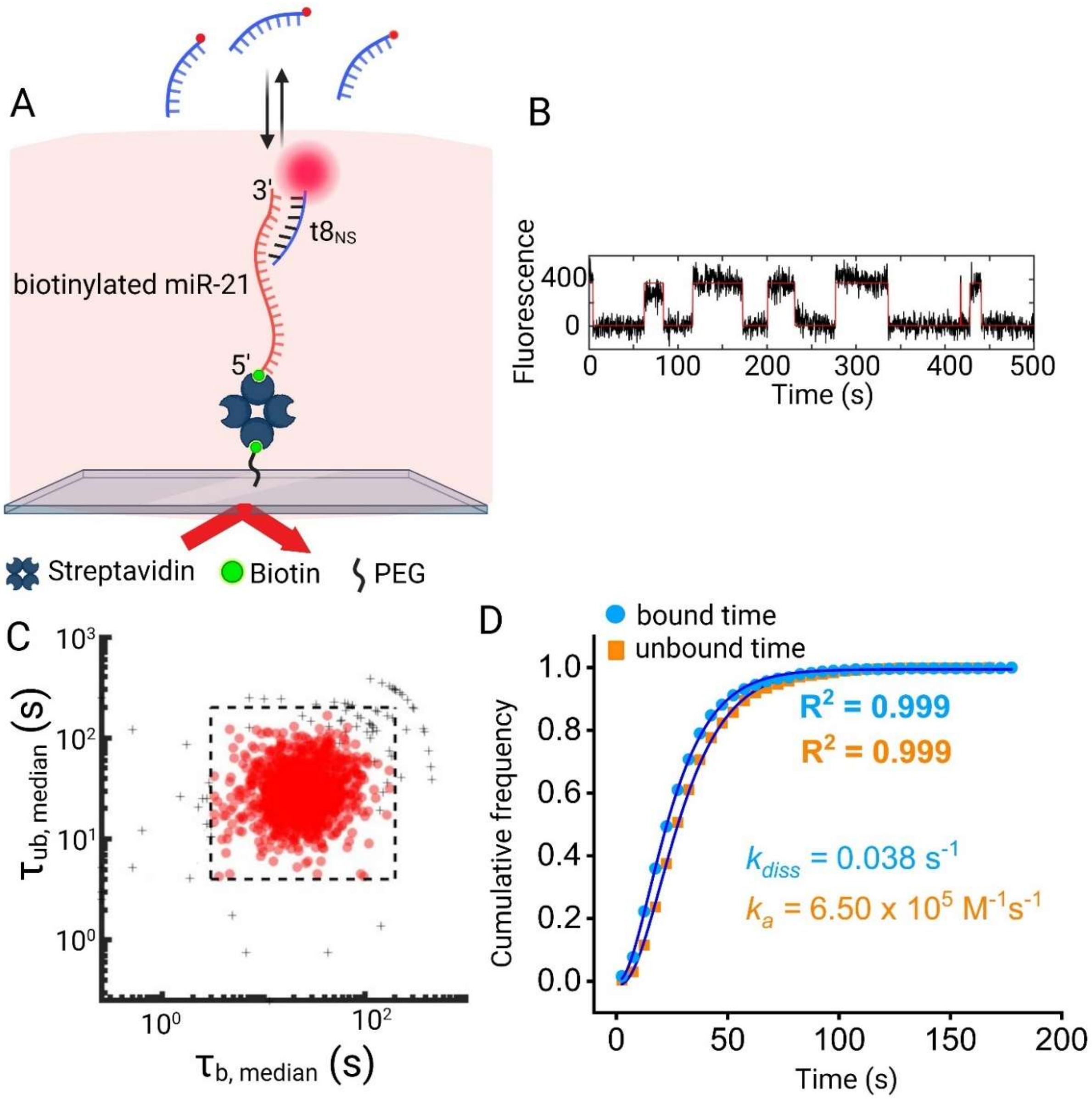


**Figure S18. Kinetics of protein-free miRNA-21 with t8$_{NS}$ probe. Related to Figure 4**

Interaction of t8$_{NS}$ probe with protein-free biotinylated miR-21. (A) Intensity vs time trajectories of interaction between t8$_{NS}$ and protein-free biotinylated miR-21. (B) Scatter plots of the median $\tau_{bound}$ and $\tau_{unbound}$ dwell times for all intensity-versus-time trajectories within a single field of view. (C) Cumulative distribution of unbound time and bound time of $t8_{NS.}$ The association and dissociation rate constants were calculated from the gamma distribution fitting of the respective cumulative distributions.

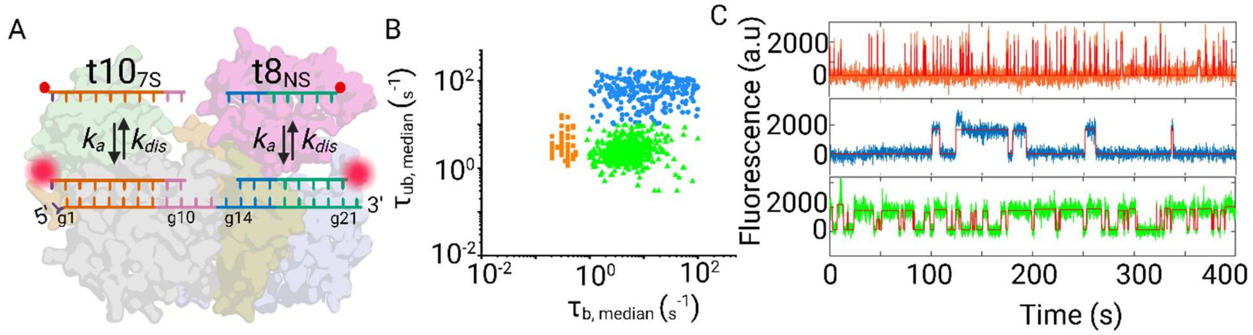


**Figure S19. Investigating the interplay between seed and non-seed binding. Related to Figure 5**

(A) Schematic showing the design of Cy5-labeled seed (t10$_{7S}$) and non-seed probes (t8$_{7S}$) employed to simultaneously investigate accessibility of seed and non-seed regions of miR-21 RISC. (B) Scatter plot of the median $\tau_{bound}$ and $\tau_{unbound}$ dwell times when both seed (t10$_{NS}$) and non-seed probes (t8$_{NS}$) are employed simultaneously. The scatter plot shows three distinct clusters whose kinetics are consistent with miRISC populations exclusively accessible to either the seed (green cluster) or non-seed (blue and orange cluster) probe. (C) Representative intensity *versus* time traces of miR-21 RISCs whose kinetics are suggestive of interaction with non-seed probe only (upper and middle traces) or seed probe only (lower trace) within the observation time window of 400 s.

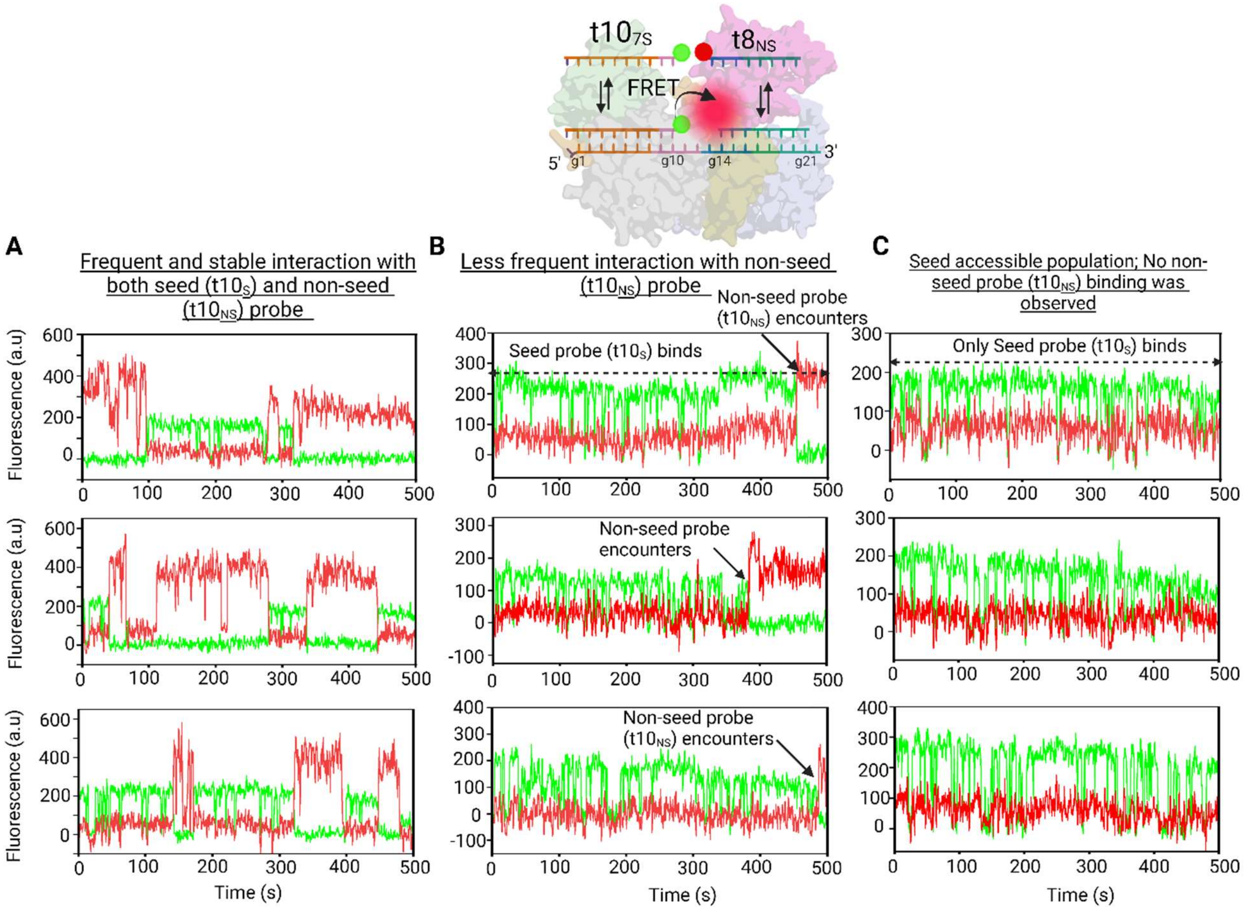


**Figure S20. Representative FRET trajectories showing simultaneous seed and long-lived supplementary plus non-seed binding**. **Related to Figure 5.**

(A) Example trace showing seed probe is bound throughout the detection window and non-seed probe frequently encounters. (B) Example trace showing seed probe is bound throughout the detection window and non-seed probe interacts very infrequently. This indicates that although seed sequence is readily accessible for binding, non-seed probe binding is conformationally restricted. (C) Example trace showing seed probe is bound throughout the detection window, but no non-seed probe binding is observed.
